# Evolution of gastrointestinal tract morphology and plasticity in cave-adapted Mexican tetra, *Astyanax mexicanus*

**DOI:** 10.1101/852814

**Authors:** Misty R. Riddle, Fleur Damen, Ariel Aspiras, Julius A. Tabin, Suzanne McGaugh, Clifford J. Tabin

**Affiliations:** Department of Genetics, Blavatnik Institute, Harvard Medical School, Boston, MA 02115; Ecology, Evolution, and Behavior, University of Minnesota, Saint Paul, MN

## Abstract

The gastrointestinal tract has evolved in numerous ways to allow animals to optimally assimilate energy from different foods. The morphology and physiology of the gut is plastic and can be greatly altered by diet in some animals. In this study, we investigated the evolution and plasticity of gastrointestinal tract morphology by comparing laboratory-raised cave- and river-adapted forms of the Mexican tetra, *Astyanax mexicanus*, reared under different dietary conditions. In the wild, river-dwelling populations (surface fish) consume plants and insects throughout the year, while cave-dwelling populations (cavefish) live in a perpetually dark environment and depend on nutrient-poor food brought in by bats or seasonal floods. We found that multiple cave populations converged on a reduced number of digestive appendages called pyloric caeca and that some cave populations have a lengthened gut while others have a shortened gut. Moreover, we identified differences in how gut morphology and proliferation respond to diet between surface fish and cavefish. Using a combination of quantitative genetic mapping, population genetics, and RNA sequencing, we found that changes to the molecular and genetic pathways that influence cell proliferation, differentiation, and immune system function may underlie evolution of the cavefish gut.

## Introduction

The gastrointestinal tract consists of functional regions distinguished by the organization of muscle and mucosal layers and cellular composition of epithelium lining the lumen. The secretory and absorptive cell types of the epithelium are constantly being replaced by self-renewing stem cells. Gut homeostasis requires integration of intrinsic signaling pathways set up during development, with external cues. For example, in the mammalian gut, Notch and WNT signaling is essential for establishing the stem cell niche and balancing stem cell renewal and differentiation of cell types (reviewed in 1). High-fat diet alters these pathways to favor self-renewal and secretory cell formation (2, 3). Cytokine signals from tissue-resident immune cells can also promote stem cell renewal in response to infection (4, 5). While studies utilizing model organisms and organoid culture have begun to reveal how homeostasis of the epithelium is maintained, there is a limited understanding of how these pathways evolve as animals adapt to different diets.

Evolutionary changes to gut morphology, such as expansion of functional domains, have been correlated to altered patterns of gene expression during development (6). The organization of the gut epithelium into crypts that support villi is important for stem cell maintenance in mammals, but in other species the epithelium is characterized by irregular folds, zig-zags, honeycomb patterns, or a spiral fold (7, 8). The presence of spatially restricted gut stem cells is common across distant phyla (i.e. arthropods and chordates) (9–11). Although it has been demonstrated that diet influences gut morphology in a number of species (reviewed in 12), evolution of the signaling pathways that integrate internal and external cues is not well understood. For animals that survive months without food, like snakes and migrating birds (13–15), altering the pathways that control homeostasis of the epithelium was likely critical to balance nutrient assimilation and energy conservation.

*Astyanax mexicanus* is a species of fish that exists as river- and cave-adapted populations that evolved on very different diets (16, 17). The river (surface) fish have access to insects and plants in abundance, while the cavefish are dependent on predominantly bat droppings and material brought in by seasonal floods (16, 17). There are a number of cave populations, named for the caves they inhabit (i.e. Tinaja, Pachón, Molino). Based on whole genome sequencing analysis, these populations reflect two independent cave derivations from surface fish within the last 200,000 years; Tinaja and Pachón cavefish populations form a separate clade from Molino cavefish (18). Cavefish have converged on similar morphological changes, such as eye loss and increased sensory structures (19–21), behavioral changes such as reduced sleep (22–25), and metabolic changes such as increased fat accumulation and starvation resistance (26–28). We previously showed that Pachón cavefish have altered gastrointestinal motility during post-larval growth to slow food transit, possibly to achieve increased nutrient absorption (29).

In this study, we explore whether cave-adaptation has led to changes in the adult gut. First, we describe the anatomy and histology of the *A. mexicanus* gastrointestinal tract, noting distinct functional regions. Second, we compare *A. mexicanus* surface and cave populations fed the same diet and discover differences in pyloric caeca number and gut length. We show that the homeostasis and morphology of the gut responds differently to changes in diet comparing Tinaja cavefish to surface fish. To investigate the genetic architecture controlling gut length, we carry out a Quantitative Trait Locus (QTL) analysis, and identify a significant QTL associated with hindgut length. We examine genes within the QTL using population genetics and RNA sequencing and reveal genetic pathways that have been altered during the evolution of gut morphology in *A. mexicanus*.

## Results

### Anatomy and histology of the adult Astyanax mexicanus gastrointestinal (GI) tract

The GI tract of teleost fish is commonly divided into four sections from anterior to posterior: head gut, foregut, midgut, and hindgut. These sections are easily defined in *A. mexicanus* and can be further subdivided based on distinct morphology and histology. The foregut consists of the esophagus, stomach, and pylorus (Figure 1A, B). The esophagus has two perpendicular layers of striated muscle and a stratified epithelium of mucus secreting cells, identifiable by a lack of staining with hematoxylin and eosin (Figure 1A). It connects to the J-shaped stomach that has three muscle layers: outermost longitudinal, the middle and thickest circumferential, and innermost oblique (Figure 1B). Gastric glands are evident in the stomach mucosa; at the base of the glands, pepsin-secreting chief cells are distinguishable by eosinophilic granules. The lumen-facing epithelium of the stomach consists of columnar mucus-secreting cells. The stomach ends at the pylorus that has thick muscle layers and connects the foregut to the midgut.

**Figure 1.**
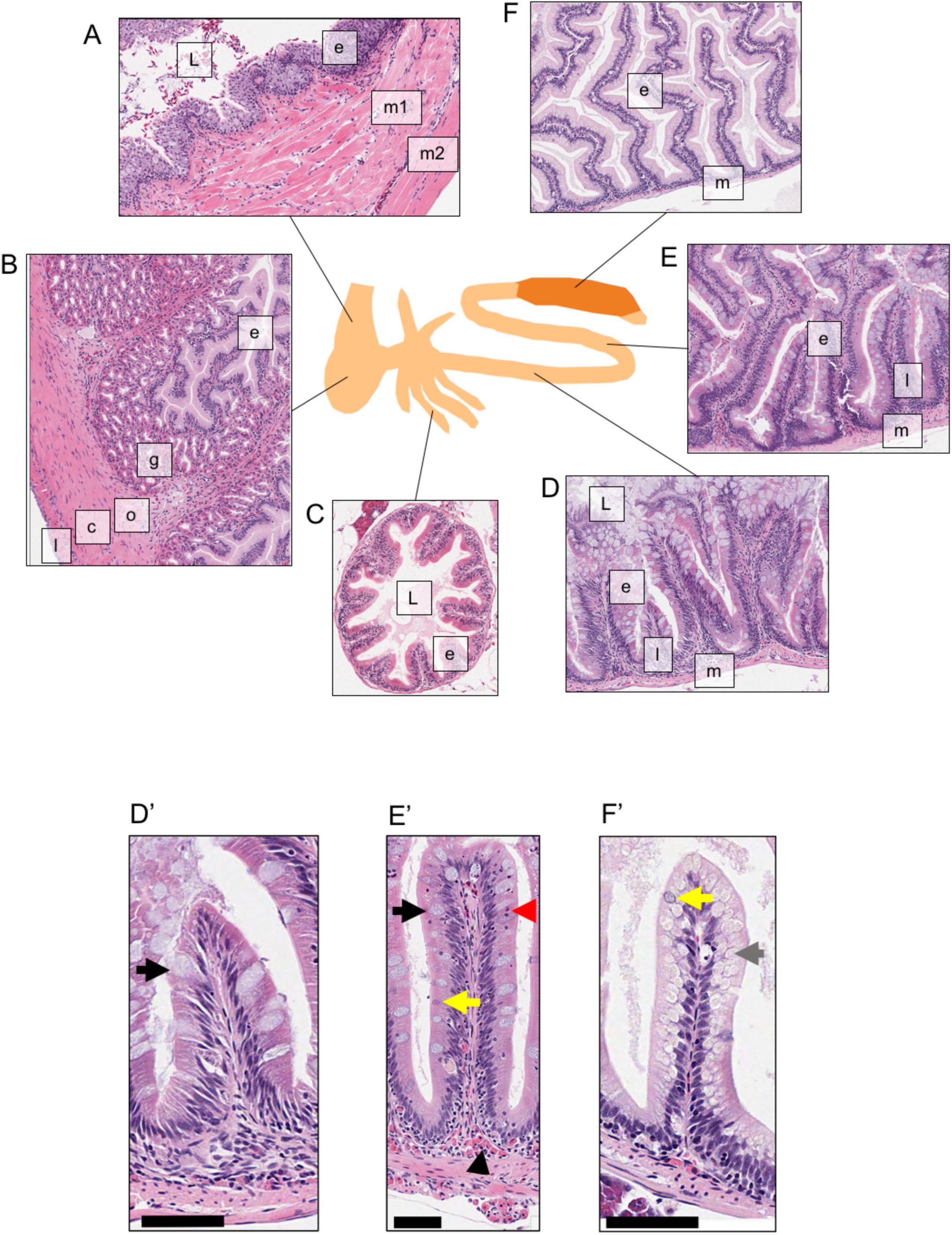
Histology of the *Astyanax mexicanus* surface fish gastrointestinal tract as shown by hematoxylin and eosin staining. A, Transverse section of esophageal region (L: lumen, e: epithelium, m1-2: muscle layers). B, Stomach section showing epithelium (e), eosinophilic gastric glands (g), and layers of muscle (o: orthogonal, c: circumferential, l: longitudinal). C, Cross section of a pyloric caecum showing the lumen (L) and ridged epithelium (e). D, Transverse section of the proximal midgut showing lumen filled with mucus (L), epithelium (e), lamina propria (l), and circumferential inner and longitudinal outer muscle layers (m). E, Transverse section of a more posterior section of the midgut showing epithelium (e), lamina propria (l), and circumferential inner and longitudinal outer muscle layers (m). F, Transverse section of the hindgut showing epithelium (e), and muscle layers (m). A-F image magnification is 10X. D-F’ 20X images (scale bar is 50µM) showing the cell types of the gut epithelium in the proximal midgut (D’), distal midgut (E’), and hindgut (F’). Black arrows highlight the differences between secretory goblet cells in the proximal (D’) and distal (E’) midgut. Black arrow head in E’ indicates group of immune cells (pink). Red arrow head shows pyknotic nuclei. Grey arrow in F’ highlights cells with hydrophobic (mostly clear) structures within the cytoplasm. D-E are images from the same slide.

The midgut is the longest portion of the GI tract. The proximal end of the midgut contains pyloric caeca: pouch structures that have a thin layer of outer connective tissue and lamina propria, and an inner epithelium organized into ridges that run from the base, to the tip of the pouch (Figure 2A). The epithelium is made up of mostly columnar enterocytes and a few mucus secreting goblet cells (Figure 1C). There are four pyloric caeca on the right-hand side of the midgut (Figure 2, “1-4”), two on the left-hand side (“5-6”), and as many as three that are shorter and more proximally restricted (“7-9”). Caeca one through five are present in fish of all four populations (surface, Tinaja, Pachón, and Molino) while six through nine are variable. Surface fish are more likely to have 1-9, Tinaja and Molino 1-6, and Pachón 1-7 (Figure 2C).

**Figure 2.**
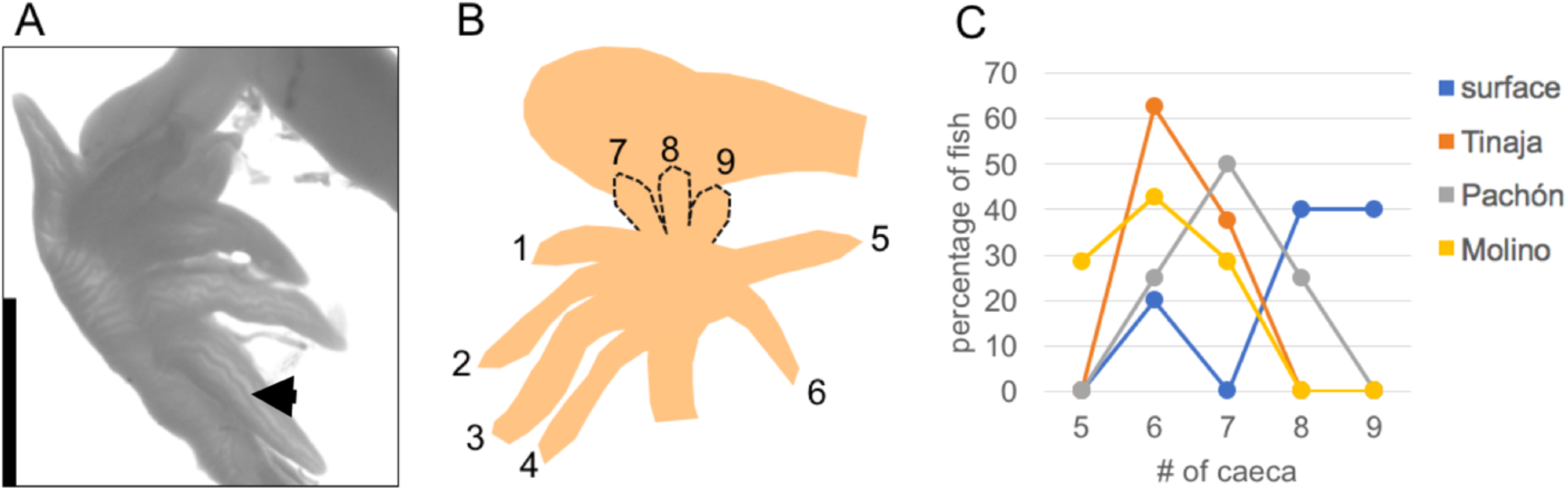
Pyloric caeca structure and number in *Astyanax mexicanus* surface and cave morphs. A, Image of the region of gut containing pyloric caeca. The darker lines show the ridged morphology of the epithelium (black arrow, scale bar is 2 mm). B, Drawing of the pyloric caeca indicating the position and number of caeca. The caeca at position 7, 8, and 9 are variable. C, Graph comparing percentage of one-month old fish with the indicated number of total caeca for each population (n=5, 6, 8, 6).

The remaining tubular portion of the midgut has inner circumferential and outer longitudinal muscle layers, and the epithelium is folded circumferentially at irregular angles. The epithelium of the proximal midgut is mostly mucus secreting goblet cells that completely lack H&E staining (Figure 1D, D’, black arrow), compared to the epithelium of the distal midgut that has goblet cells with basophilic granules (Figure 1E’). The lumen of the distal midgut also contains basophilic enteroendocrine cells that are distinguishable by a smaller size and more apical location (Figure 1E’ yellow arrow). A greater number of eosinophilic immune cells are also evident in the lamina propria of the distal region of the midgut (Figure 1E’ black arrowhead).

The proximal hindgut is wider than the midgut and has thinner muscle and lamina propria layers (Figure 1F-F’). The enterocytes in the epithelium are highly vacuolated likely representing absorbed hydrophobic molecules (Figure 1F’, grey arrow). Apically located enteroendocrine cells are also evident in the epithelium (Figure 1F’, yellow arrow). The distal hindgut (rectum) has a thicker muscle wall and connects the GI tract to the anus. Overall, the organization of the *A. mexicanus* gut is similar to other Characiformes. Having established the general anatomy and histology of the GI tract using the surface form, we were next able to explore differences between the surface and cave populations.

### Ecotypic differences in gut length

The length of each region of the GI tract is an important variable determining digestive efficiency and can be altered by diet. To determine the ecotypic differences in gut length between populations, we compared relative gut length (gut length divided by fish length) between individually housed fish fed the same diet (38% protein, 7% fat, 5% fiber) and amount per day (6mg). We found that Tinaja (n=6) and Molino (n=5) cavefish tend to have a longer midgut, and Pachón (n=6) cavefish tend to have a shorter midgut compared to surface fish (n=6) (Figure 3A, B). Tinaja and Molino also tend to have a longer hindgut and Pachón have a significantly shorter hindgut compared to surface fish (Figure 2C, p=0.05, one-way ANOVA with Tukey’s post hoc test). Interestingly, while all fish were dissected 24 hours post-feeding, we noted a difference in the amount of fecal matter in the gut (Figure 3A). All of the Tinaja cavefish guts were entirely full (n=6), compared to one out of six Pachón, and four out of five Molino. All of the surface fish guts were mostly empty as seen in Figure 3A (n=6). In summary, there are moderate differences in adult gut length between populations that are independent of diet, and some adult cavefish populations may have slower gastrointestinal transit.

**Figure 3.**
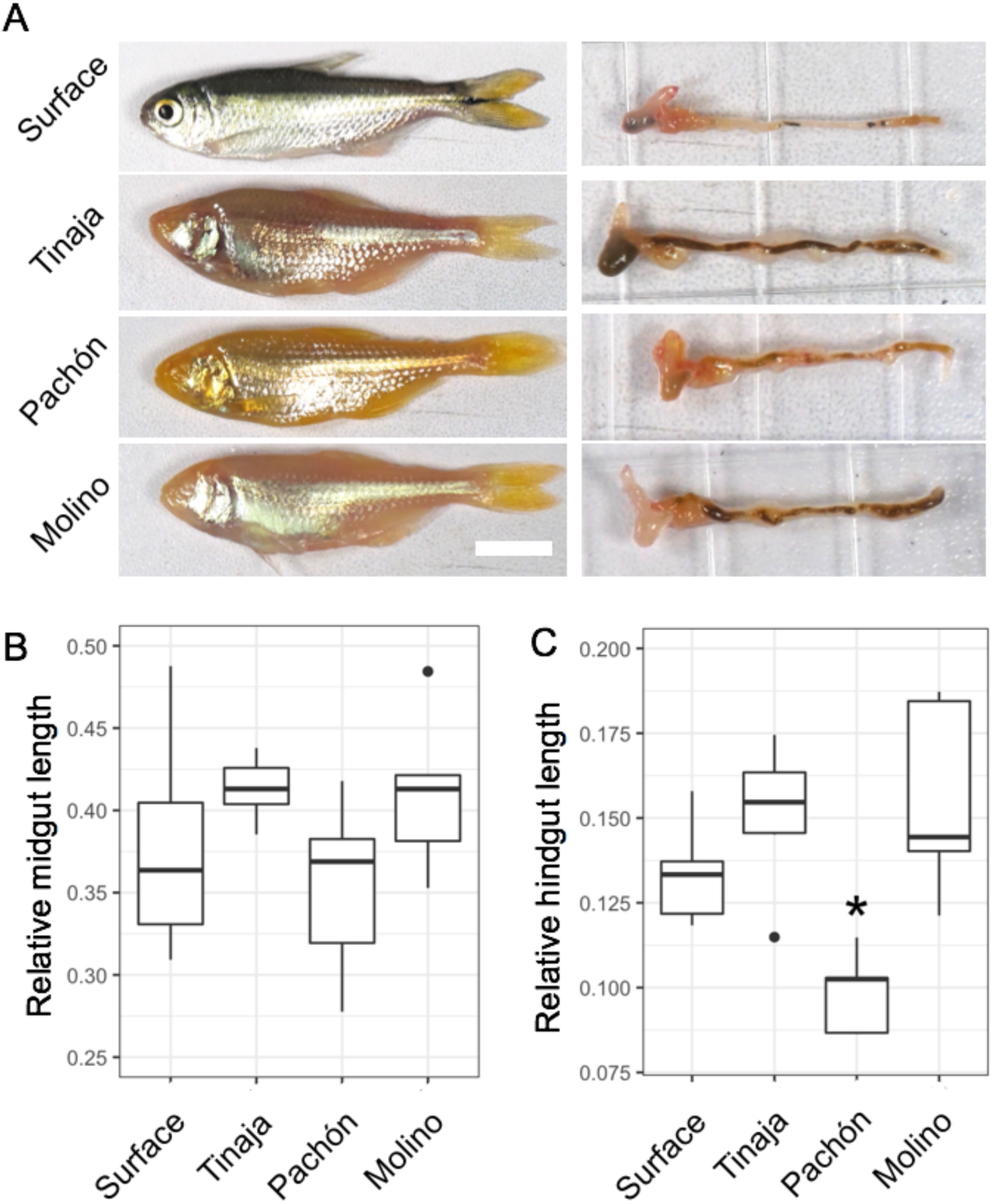
Relative length of the gastrointestinal (GI) tract of *Astyanax mexicanus* surface and cave morphs. A, Images of fish from the indicated populations that were fed 6 mg of pellet food per day for greater than 8 months and their GI tracts (scale bars is 1cm). B, Boxplots showing relative midgut and hindgut length of fish from the indicated populations (n=6 for surface, Tinaja, and Pachón. n=5 for Molino). For box plots, median, 25th, 50th, and 75th percentiles are represented by horizontal bars and vertical bars represent 1.5 interquartile ranges. Asterisks indicates Pachón relative hindgut length is significantly less than surface (p=0.05), Tinaja (p=0.002), and Molino (p=0.001, one-way ANOVA with Turkey’s post hoc test).

### Diet influences gut morphology differently in surface fish and Tinaja cavefish

Surface-adapted *A. mexicanus* consume plants and insects throughout the year. In contrast, cavefish populations are subject to seasonal fluctuations, including periods of deprivation. We hypothesized that cavefish may have evolved increased plasticity in gut morphology to maximize nutrient absorption during times of abundance and conserve energy when food is scarce. To test this hypothesis, we switched the diet of adult surface fish and Tinaja cavefish from moderate nutrient content (7% fat, 38% protein, 5% fiber) to either low-nutrient (4% fat, 32% protein, 3% fiber) or high-nutrient (18% fat, 55% protein, 2% fiber). After eight months, we measured the length of the gut segments, as well as the circumference and average number of folds in cross sections, and compared the values to fish that remained on the moderate nutrient diet (Figure 4, n=3 fish per population and diet, and 3 histological sections per segment). On the moderate nutrient diet fed *ad libitum*, Tinaja tend to have a shorter midgut that is wider and has more folds compared to surface fish (relative midgut length: surface = 0.45, Tinaja = .36, circumference: surface = 0.16cm, Tinaja = 0.20, circumference normalized to fish length: surface = 0.03, Tinaja = 0.04, fold number: surface = 6, Tinaja = 9, Figure 4 A-C). Tinaja also tend to have a longer hindgut on this diet as was observed on the controlled diet (surface = 0.17, Tinaja = 0.20).

**Figure 4.**
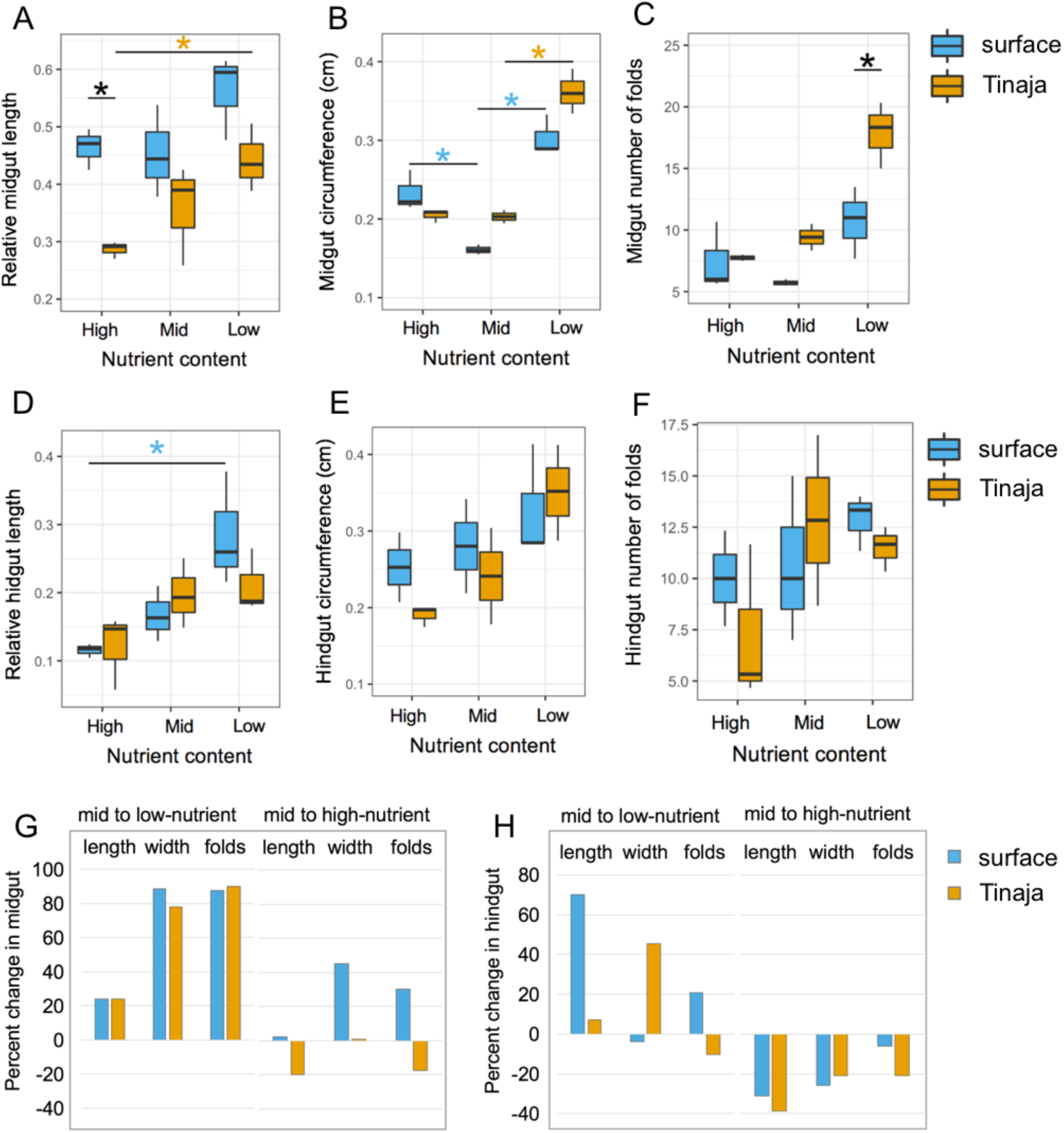
Diet influences *A. mexicanus* gut morphology. Relative length, circumference, and number of folds in the epithelium in the midgut (A-C) and hindgut (D-E) in fish continuously fed a moderate diet or switched to a high-nutrient or low nutrient diet as adults (n=3 fish per population and diet, and 3 tissue sections average per individual fish). In boxplots, median, 25th, 50th, and 75th percentiles are represented by horizontal bars and vertical bars represent 1.5 interquartile ranges. Significance code (*p<0.05) from one-way ANOVA with HSD post hoc test comparing between populations (black asterisks) or within population by diet (orange asterisks = Tinaja, blue asterisks = surface fish). Bar graphs show percent change in the indicated phenotype in the midgut (G) and hindgut (H).

In fish that were switched to a low-nutrient diet, we observed a 24% increase in the length of the midgut in both populations (surface = 0.45 to 0.56, p=0.36; Tinaja = 0.36 to 0.44, p = 0.60; one-way ANOVA with HSD post-hoc test, Figure 4A, G). The circumference of the midgut also increased significantly by 89% in surface fish (0.16 to 0.30 cm, normalized to fish length 0.03 to 0.06, p<0.005) and 78% in Tinaja cavefish (0.20 to 0.36 cm, normalized to fish length 0.04 to 0.07, p<0.005, Figure 4B, G). The number of folds in the midgut epithelium increased 2-fold in Tinaja (9 to 18) and nearly 2-fold in surface fish (6 to 11). The hindgut length increased by 70% in surface fish (0.17 to 0.28) and only 7% in Tinaja cavefish (0.20 to 0.21, Figure 4D, H). The circumference of the surface fish hindgut did not change significantly (0.34 to 0.33cm, normalized to fish length = 0.06 to 0.07) and the circumference of the Tinaja hindgut increased by 45% in Tinaja (0.24 to 0.35cm, normalized to fish length =0.04 to 0.07, Figure 4D, H). The number of folds in the hindgut epithelium changed only slightly in both populations (Tinaja = 13 to 12, surface = 11 to 13). In summary, midgut length, width, and fold number increased similarly in both populations in response to a low-nutrient diet (Figure 4G). In contrast, the hindgut responds by growing in length in surface fish and, growing in width in cavefish.

In fish switched from a moderate to a high-nutrient diet, the length of the midgut did not change in surface fish (0.45 to 0.46, Figure 4A) and decreased in Tinaja cavefish by 20% (0.36 to 0.29). The circumference of the midgut increased in surface fish (0.16 to 0.23cm, normalized to fish length = 0.03 to 0.05) and did not change in cavefish (0.20 to 0.20cm, normalized to fish length = 0.04 to 0.04; Figure 4C). The average number of epithelial folds increased slightly in surface fish (6 to 7) and decreased slightly in the cavefish (9 to 8, Figure 4C). Normalized hindgut length decreased in both populations in response to a high-nutrient diet, although the decrease was greater in Tinaja: 39% (0.20 to 0.12) versus 31% in Surface fish (0.17 to 0.12, Figure 4B). The circumference of the hindgut also decreased, by 26% in surface fish (0.34 to 0.25cm, normalized to fish length = 0.06 to 0.05) and by 21% in cavefish (0.24 to 0.19cm) although the value normalized to fish length did not change in cavefish (0.04 to 0.04, Figure 4D). The number of folds in the hindgut decreased by one in surface fish (11 to 10) but more strikingly in Tinaja cavefish reduced to almost half (13 to 7, Figure 4C). In summary, the Tinaja cavefish gut is more responsive than surface fish to a high-nutrient diet since Tinaja cavefish exhibit a greater reduction in midgut and hindgut length and hindgut fold number. Overall, the results suggest that plasticity of the gut is dependent on the type of diet and region of the gut and reveal fundamental differences between the populations.

### Differences in the regulation of gut proliferation between surface fish and Tinaja cavefish

The gastrointestinal epithelium undergoes constant turnover, being replenished by populations of dividing cells that are restricted to the base of villi or epithelial folds (1). We next sought to understand how differences in morphology are achieved by investigating homeostasis of the gut epithelium. We injected fish on the low-nutrient diet with a thymidine analog (5-ethynyl-2’-deoxyuridine, EdU) and, after 24 hours, determined the number of EdU positive cells as a measure of the amount of cell division (Figure 5A, B, F, J). We found that Tinaja cavefish had on average over four times as many EdU positive cells in the midgut (Figure 5F, surface = 103, Tinaja = 449, p = 0.063 two-tailed t-test). Higher proliferation corresponds to a significantly greater number of folds in the midgut epithelium in Tinaja cavefish (Figure 4C, average surface = 11, Tinaja = 18; p = 0.04, two-tailed t-test). However, the number of EdU positive cells per fold in the midgut is also greater in Tinaja (average surface = 11, Tinaja = 26; p = 0.09); EdU positive cells in the epithelium appear more restricted to the base of the folds in surface fish, whereas they extend further up the fold in Tinaja cavefish (Figure 5A, B). In the hindgut, the average number of EdU positive cells is also greater in Tinaja (Figure 5J, surface = 154, Tinaja = 354; p = 0.17), although there is only a slight difference in the number of folds (surface = 13, Tinaja = 12). These results suggest that altered homeostasis of the gut epithelium could drive morphological differences between the populations.

**Figure 5.**
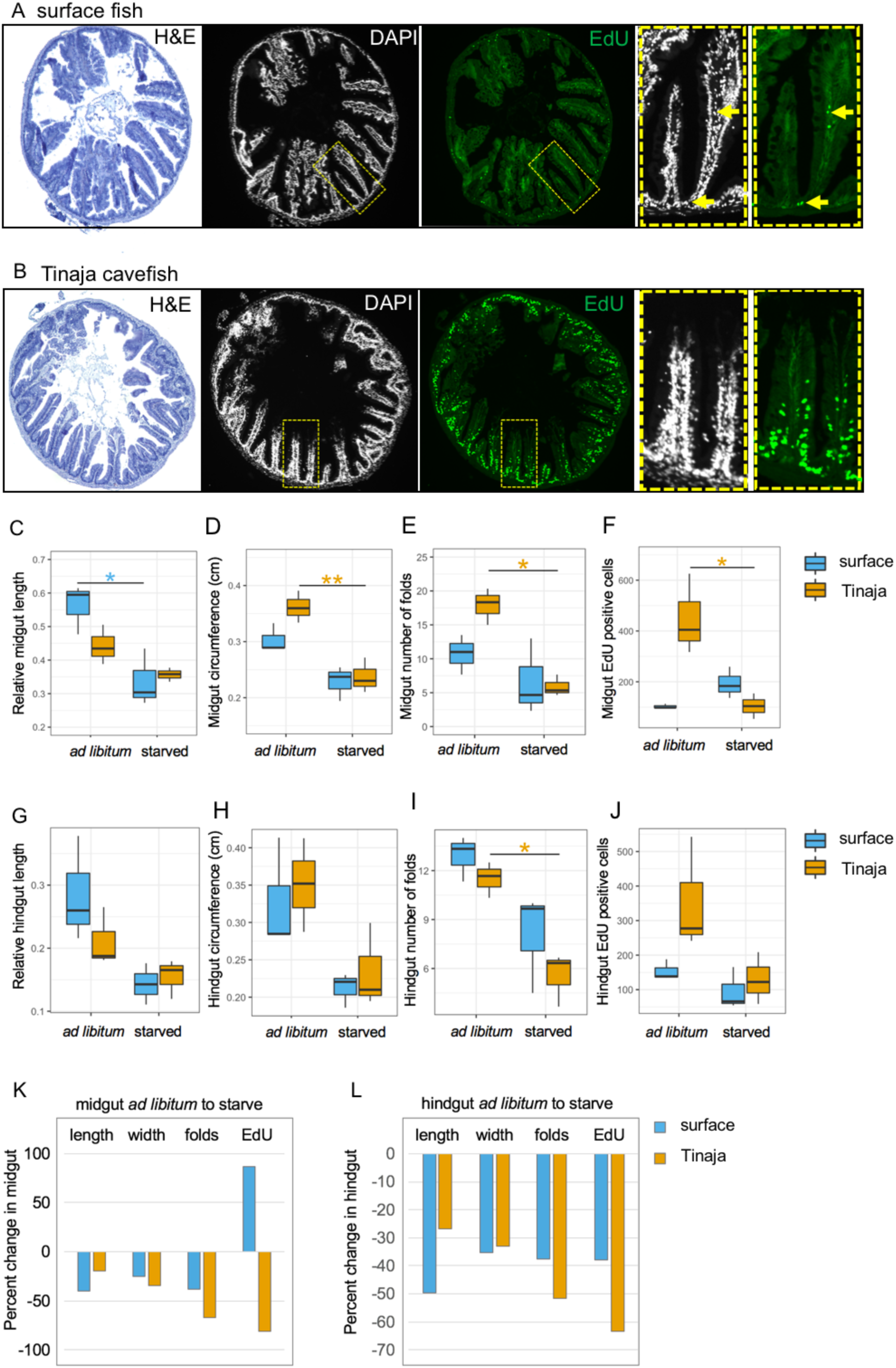
Cavefish have more proliferation in the intestinal epithelium on a low-nutrient diet compared to surface fish and, reduce proliferation in response to starvation. Cross section of midgut in surface fish (A) and Tinaja cavefish (B) stained with H&E (left) and serial section showing DAPI staining and EdU positive cells; larger view of yellow boxed region shown in right panels. Yellow arrows point to EdU positive cells. Note yellow arrow near the tip of the fold shows cell in the lamina propria. C-F, Quantification of midgut length, circumference, number of folds, and EdU positive cells in fish fed *ad libitum* versus starved for two weeks. G-J, Quantification of hindgut length, circumference, number of folds, and EdU positive cells in fish fed *ad libitum* versus fasted for two weeks (n=3 fish per population and diet and 3 tissue sections averaged per individual fish). In boxplots, median, 25th, 50th, and 75th percentiles are represented by horizontal bars and vertical bars represent 1.5 interquartile ranges. Significance code (*p<0.05, **p<.005) from one-way ANOVA with HSD post hoc test. K-L, Bar graphs showing percent change in the indicated phenotypes in the midgut (K) and hindgut (L).

Cavefish evolved in a nutrient-limited environment and unlike surface fish are subject to periods of starvation. Because cell-turnover is energetically expensive, we hypothesized that cavefish may have evolved mechanisms to limit the proliferation in the gut when food is not available. To test this, we starved fish (for a period of two-weeks) that were previously fed the low-nutrient diet, and compared EdU incorporation over a 24-hour period (Figure 5F, J, K, L). The number of EdU positive cells in the Tinaja midgut decreased significantly by 81% (average 449 fed versus 85 starved, p = 0.005, one-way ANOVA comparing diet and population). In contrast, the number of EdU positive cells nearly doubled in surface fish (average 103 fed versus 193 fasted, p = 0.63). In both populations, midgut length, circumference, and fold number decreased in response to starvation (Figure 5C-D). In surface fish, the midgut length shortened significantly by 40% (0.56 to 0.34, p=0.01, one-way ANOVA) compared to only 19% in Tinaja cavefish (0.44 to 0.36, p=0.42). The midgut circumference decreased by 25% in surface fish (0.30 to 0.23cm, normalized to fish length = 0.06 to 0.05) and significantly in cavefish by 34% (0.36 to 0.24cm, p=0.004, normalized to fish length = 0.07 to 0.05; p=0.03). The number of folds in the midgut decreased from 11 to 7 in surface fish (p=0.53) and more dramatically, from 18 to 6 in Tinaja cavefish (p=0.01). In summary, Tinaja cavefish reduce proliferation in response to starvation and this is associated with a substantial decrease in midgut circumference and fold number, but a more moderate decrease in length, compared to surface fish.

In the hindgut, the average number of EdU positive cells also reduced significantly in Tinaja cavefish in response to starvation (Figure 5J, average 354 to 130, p=0.005, one-way ANOVA), and to a lesser extent in surface fish (38%, average 154 to 95, p= 0.63 one-way ANOVA). Relative hindgut length reduced by 50% in surface fish (0.28 to 0.14) and only 20% in Tinaja (0.21 to 0.15). We observed a similar reduction in hindgut circumference, by 35% in surface fish (0.33 to 0.21cm, normalized to fish length = 0.07 to 0.04) and 34% in Tinaja cavefish (0.35 to 0.23cm, normalized to fish length = 0.07 to 0.05). The number of hindgut folds reduced from 13 to 8 in surface fish and 12 to 6 in Tinaja cavefish. Similar to what occurs in the midgut, in the hindgut, Tinaja decrease proliferation more substantially in response to starvation but have a less drastic decrease in length compared to surface fish. Combined, our findings support the hypothesis that cavefish have mechanisms for tuning gut homeostasis in response to food intake that differ from surface fish.

### Genetic mapping reveals quantitative trait loci associated with hindgut length

We next sought to understand the genetic basis of differences in gut morphology between surface fish and cavefish. We carried out a quantitative trait loci (QTL) analysis using F2 surface/Tinaja hybrids (see methods). To eliminate the effect of diet or appetite, we individually housed the hybrids as adults and ensured they consumed the same diet (38% protein, 7% fat, 5% fiber) and amount per day (6mg). After four months, we determined the length and weight of the fish, number of pyloric caeca, and length of the midgut and hindgut. We identified a QTL for female body condition (fish weight/fish length) on linkage group 13 with a LOD score of 5.53 accounting for 21% of the variance in this trait (Figure 6B). We did not identify a QTL for male body condition (fish weight/fish length Figure 6A). We also did not identify a significant QTL for pyloric caeca number but observed a peak on linkage group 24 (Figure 6C).

**Figure 6.**
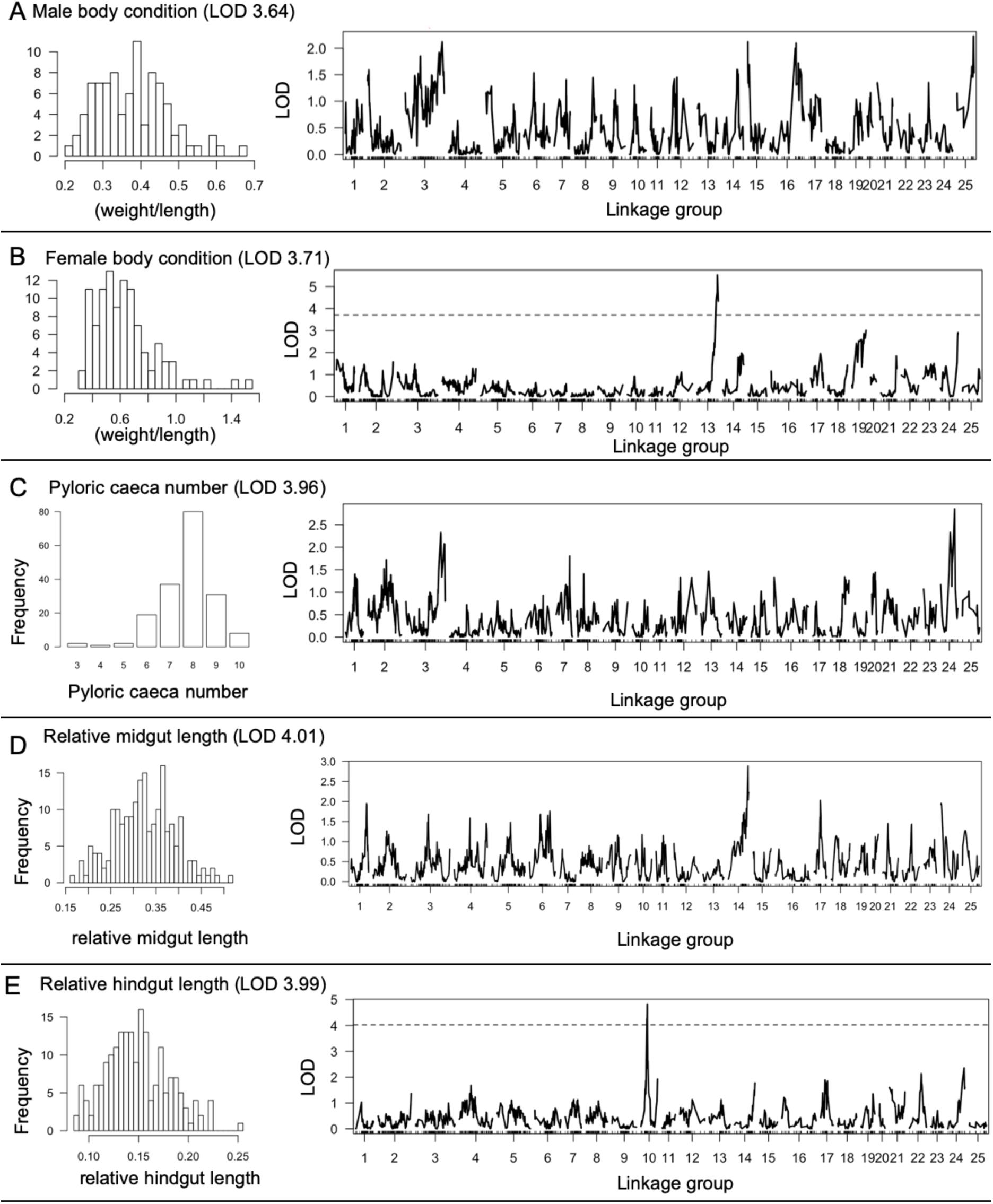
Genetic mapping of body condition and gut morphology in adult *A. mexicanus* surface/Tinaja F2 hybrids. A-E, Histograms showing distribution of the indicated phenotypes in F2 hybrids and (right) genome wide scan showing logarithm of the odds (LOD) score for the indicated phenotypes using Haley-Knott regression mapping. A, B non-parametric model, C-E normal model. Significance thresholds at p=0.05 are indicated in parenthesis following the phenotype.

There is a strong positive correlation between fish length and gut length as anticipated (midgut p-value=2.2×10^-16, hindgut p-value=5.2×10^-14, Pearson’s product-moment correlation). We therefore performed the QTL analysis using relative gut length. We did not identify a significant QTL for relative midgut length but observed a peak on linkage group 14 (Figure 6D). We found a significant QTL for relative hindgut length on linkage group 10 with a LOD score of 4.82 at the peak marker r172341 accounting for 11% of the variance in this trait. Interestingly, it is the heterozygous genotype at this position that is associated with longest relative hindgut length (Figure 7C). Similarly, F1 hybrids have a longer hindgut than surface fish or Tinaja cavefish (Figure 7A,B). The midgut of F1 hybrids is also longer; however, when we examined the effect plot for the marker with the highest LOD score for relative midgut length, we found that individuals with cave genotype have the longest midgut (Figure 7D, E). Combined, the results indicate overdominance or positive epistasis in the genetic network controlling gut length.

**Figure 7.**
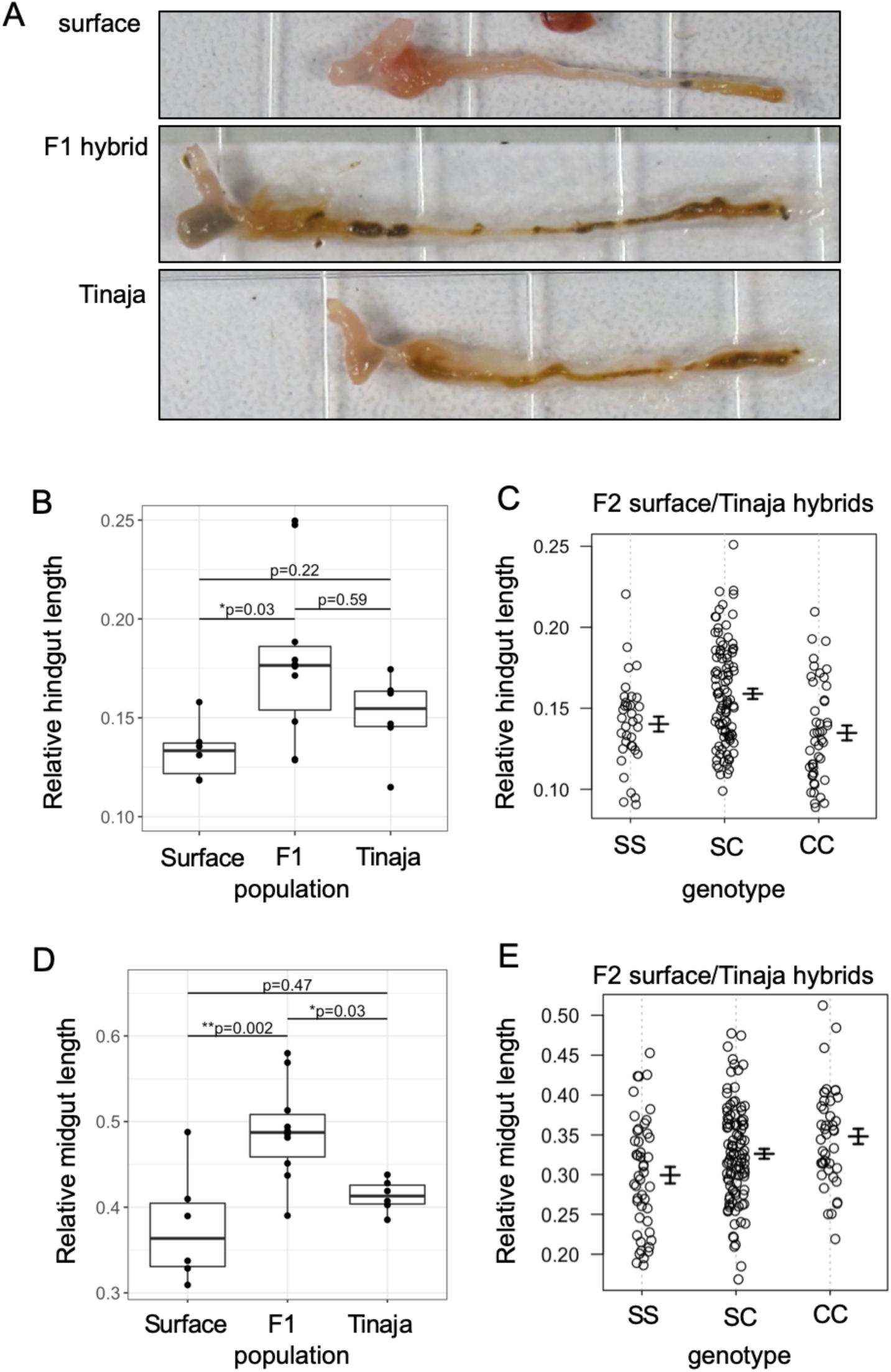
Gut length heterosis in surface/Tinaja hybrids. A, Image of gut from surface fish, surface/Tinaja F1 hybrid, and Tinaja cavefish of similar lengths. B, Quantification of relative hindgut length in surface (n=6), F1 hybrid (n=10), Tinaja (n=6). C, Quantification of relative hindgut length of surface/Tinaja F2 hybrids with the indicated genotype at the marker with the highest LOD score (SS: homozygous surface, SC: heterozygous, CC: homozygous cave). D, Quantification of relative midgut length in in surface (n=6), F1 hybrid (n=10), Tinaja (n=6). E, Quantification of relative midgut length of surface/Tinaja F2 hybrids with the indicated genotype at the marker with the highest LOD score (SS: homozygous surface, SC: heterozygous CC: homozygous cave).

### Candidate genes controlling hindgut length

To search for genetic changes that contribute to variation in hindgut length, we used a combination of population genetics and differential gene expression analysis. We first used the Pachón cavefish genome assembly to identify the position of markers that define the 1.5-LOD support interval for the hindgut QTL. We found that two markers are on scaffold KB882083 and one is on KB871783 (Table 1). We determined several population genetic metrics for all of the genes on each scaffold using whole genome sequencing data (Dataset S1, 18). This data set includes Tinaja, Pachón and Molino cave populations and two surface populations: Río Choy, which is representative of our lab strains and is more closely related to the stock of fish that invaded the Molino cave, and Rascón, which is more representative of a separate stock of surface fish that invaded the Tinaja and Pachón caves. We found that 45 genes within the 1.5-LOD support interval for the hindgut QTL have fixed differences in the coding regions between Río Choy surface fish (n=9) and Tinaja cavefish (n=10) populations (Dataset S1, all on scaffold KB882083). Ten of these genes may be under selection since the haplotype including the genes exhibits evidence of non-neutral evolution comparing Río Choy to Tinaja (Max pairwise divergence = 1 and significant HapFLK p-values (< 0.05), Table 2, Figure 8A). Two of the ten genes also fit within the same statistical parameters comparing Rascón surface fish to Tinaja cavefish (Table 2). These two genes are predicted to encode complement factor B-like proteins that are involved in innate immunity (herein referred to as Cfb1 and Cfb2). We found that *cfb1/2* also have significant HapFLK p-values comparing Pachón to surface populations, but not to Molino (Table 2). The ten genes with significant HapFLK values in Tinaja have varying ranks of divergence in the other cave populations (Figure 8A,B). Genes that show high levels of divergence in all populations may be generally important for cave adaptation (e.g. *cfb1/2*, Figure 8A, B). Genes that show divergence in only the Tinaja population (e.g. *klf5l*) may be important for adaptation only in the Tinaja cave.

**Table 1.** Location of QTL markers defining confidence internal for relative hindgut length.

| **marker** | **LOD** | **LG: position** | **Scaffold: position** | **Chromosome: position** |
| --- | --- | --- | --- | --- |
| r70838 | 1.64 | 10: 61.11 | KB871783: 144963 | 14: 27137229 |
| r172341 | 4.82 | 10: 68.28 | KB882083: 490666 | 14: 17061465 |
| r171715 | 2.82 | 10: 70.29 | KB882083: 2674927 | 14: 11724563 |

Genome-wide maximum penalized LOD score using 1000 random permutations, position on the linkage group (LG), Pachón genome scaffold, and surface fish chromosome.

**Table 2.**
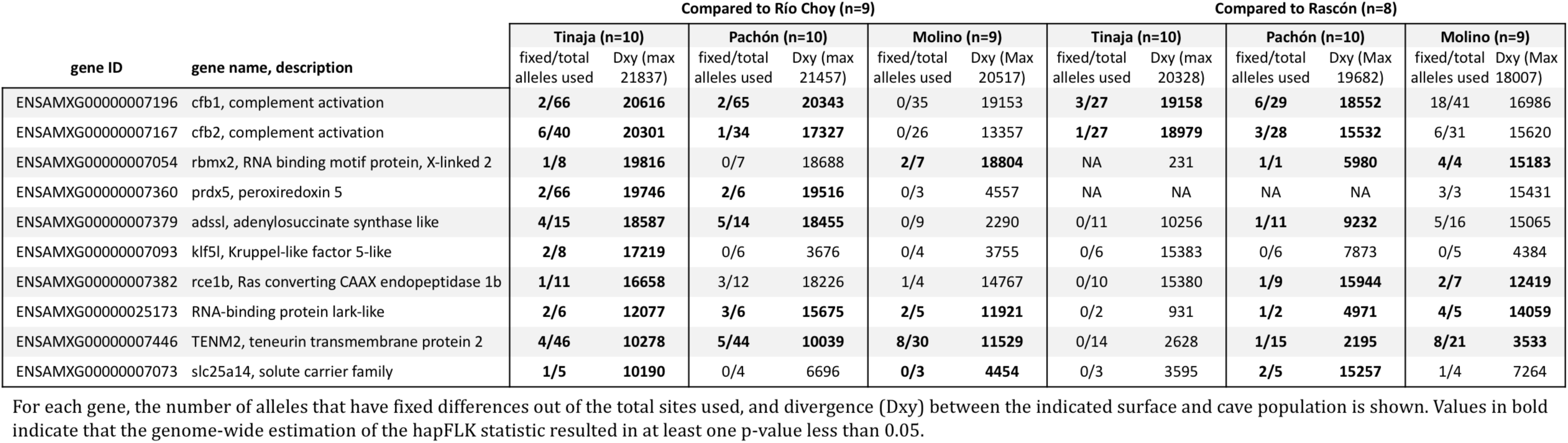
Population metrics for genes in the hindgut QTL that exhibit evidence of non-neutral evolution comparing Río Choy with Tinaja.

**Figure 8.**
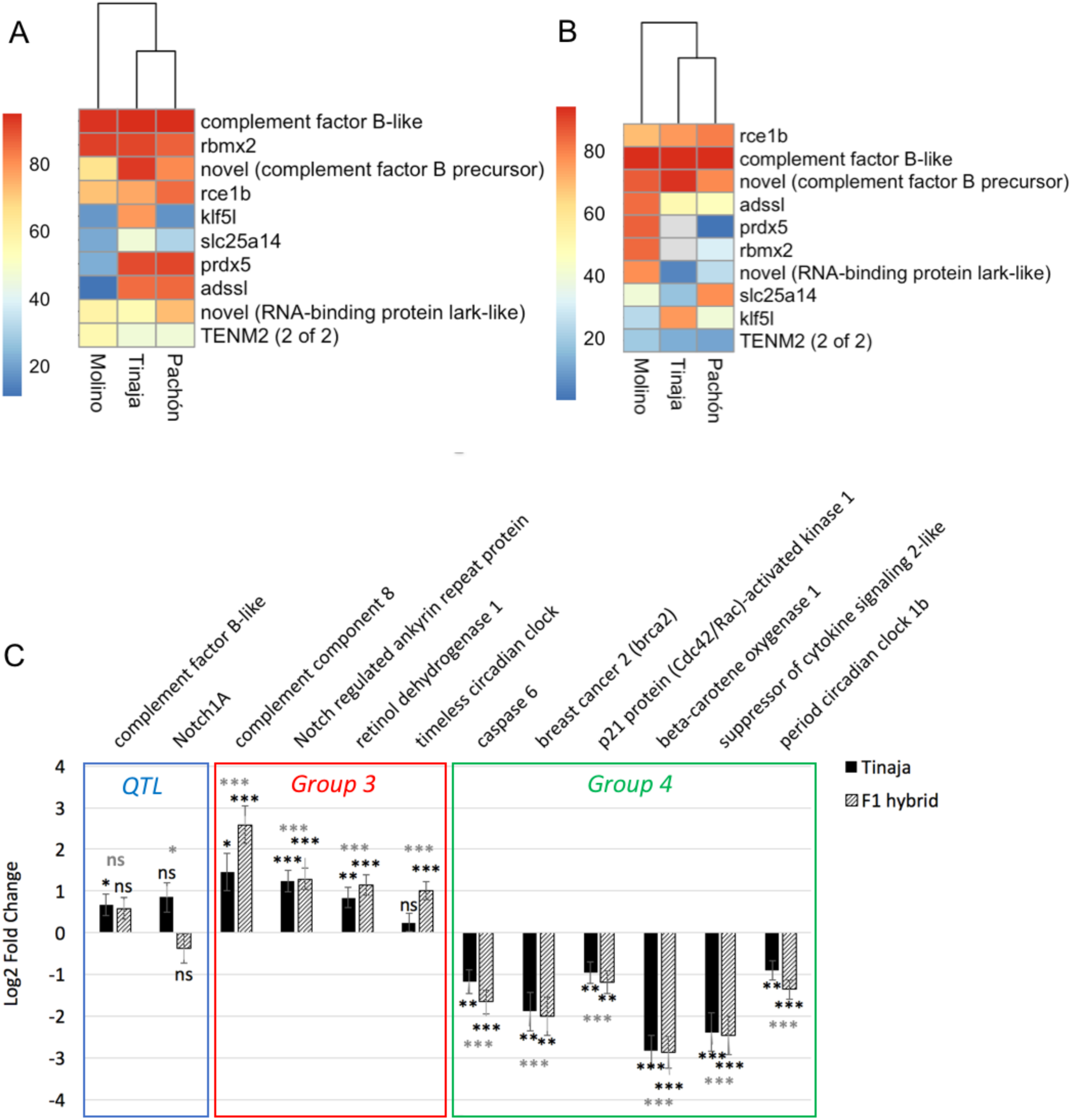
Population genetics and RNA sequencing analysis of candidate genes controlling hindgut length. A, B, Heat maps showing the genome-wide divergence percentile of genes within the hindgut QTL that have significant HapFLK p-values. A, Divergence of Río Choy surface fish from three cavefish populations, and B, Divergence of Rascón surface fish from three cavefish populations. C, Log2 Fold expression change compared to surface fish of a subset of genes within the hindgut QTL (blue), or that are a part of the gene cluster that shows lowest (group3, red) and highest (group4, green) expression in F1 hybrids. Grey significance codes from likelihood ratio test comparing all three sample types and black asterisks next to bars indicate significance from Wald test comparing to only surface fish (Significance code for adjusted p-value: *<.05 **<.005 ***<.0005, ns>.05). Error bars indicate standard error estimate for log2 fold change.

Next, we explored which genes in the QTL may have regulatory mutations by comparing levels of expression. For this analysis, we utilized the surface fish genome assembly that is organized into chromosomes. We found that the markers that define the estimated confidence interval for the hindgut QTL are on chromosome 14 and span a region of approximately 15KB (Table 1). This region has 342 genes. We determined if any of these genes are differentially expressed in the hindgut using RNA sequencing data from adult surface, Tinaja, and surface/Tinaja F1 hindguts (30) (n=5 hindguts per population). We found that 31 genes are differentially expressed using a likelihood ratio test to compare all three sample types (Table S1). Pairwise comparison between Tinaja cavefish and surface fish revealed 9 additional differentially expressed genes (Table S1). Included in this list is *cfb1*. Tinaja cavefish exhibit the highest expression of *cfb1* and expression in the F1 hybrid is intermediate (Figure 8C). We reasoned that genes showing either the greatest or lowest expression in F1 hybrids may be strong candidates for controlling differences in length since the heterozygous genotype at the QTL is associated with the longest hindgut. Of the genes that are in the QTL and differentially expressed, five show the highest expression in the hybrid and eight show the lowest expression in the hybrid. Among the genes that are lowest in the hybrid is *Notch1A* (Figure 8C, Table S1). Notch signaling controls stem cell renewal and formation of secretory versus absorptive cell fate in the intestine (31).

### Expression patterns associated with differences in gut length

We next broadened our analysis of the RNA sequencing data to investigate other pathways that may have been altered during the evolution of the cavefish gut; 3550 genes are differentially expressed between surface, F1 hybrids, and Tinaja hindguts (30). We searched for patterns across the differentially expressed genes between sample groups (degPatterns function in DEseq 2, Figure S1) and identified clusters of genes that show either greatest (Table S2, group 3, n=55) or lowest (Table S3, group4, n=53) expression in the F1 hybrid, reasoning that these may be more likely to be associated with differences in hindgut length. The genes that have the greatest expression in the hybrid include an additional component of the complement system (*C8G*), a negative regulator of Notch signaling (*nrarpa*), retinoic acid signaling components (*Cyp26A1, rdh1*), and a circadian rhythm gene (*timeless*) (Figure 8C). The genes that show a pattern of lowest expression in the hybrid include tumor suppressor and apoptosis related genes (*casp6, brca2, pak1*), a suppressor of cytokine activity (not annotated), regulator of retinoic acid production (*bco1*), and a circadian rhythm gene that typically correlates with timeless expression (*per1b*) (Figure 8C). In summary, we found that surface fish and cavefish have differences in the genetic architecture controlling hindgut length, and that cavefish have altered expression of genes controlling cell proliferation, cell signaling, and immune system function.

## Discussion

A major challenge for *Astyanax mexicanus* when they invaded caves was adapting to the drastically limited availability of nutrients compared to what they experienced in the ancestral river environment. In response, they evolved hyperphagia, starvation resistance, and increased fat accumulation (27, 28, 32). Here, we investigated how the morphology and homeostasis of their gastrointestinal tract adapted as a consequence of nutrient restriction. Our data suggest that specializations for individual caves may have occurred. By comparing fish on the same diet in the laboratory, we found that Tinaja and Molino cave populations converged on a lengthened gut while Pachón have a shortened gut. A plausible explanation for this is differences in cave ecology; the Pachón cave is less impacted by seasonal flooding and has cave-adapted micro-crustaceans that serve as a food source for post-larval fish (16). In all cave populations, we observed a reduction in the number of pyloric caeca, specifically absence of caeca 7-9. The pyloric caeca may have roles in digestion, absorption, and immune system function (33, 34). It is possible that caeca 7-9 are important for digestion of foods, or response to pathogens, that are only present in the surface fish environment, and these structures became vestigial and eventually disappeared in the cave. Other groups of fish like salmonids can have hundreds of pyloric caeca (35). Our study establishes a genetically accessible model to understand pyloric caeca function, development, and evolution.

We hypothesized that cavefish evolved increased plasticity of the gut as an adaptation to save energy during seasonal fluctuation of food. The most striking difference that we observed between populations is that cavefish have much greater proliferation in the gut epithelium compared to surface fish under fed conditions and reduce proliferation in response to food deprivation. Cavefish may have increased sensitivity in the pathways that sense or respond to nutrient shortage and it is plausible that this provides an advantage in the cave. On a high-nutrient diet, cavefish exhibit a more dramatic reduction in gut length compared to surface fish. Reducing length when nutrients are abundant could achieve a more optimal balance between energy extraction and storage; the gut is packaged within the body cavity and its size could limit the amount of visceral fat that can accumulate. How the cellular dynamics of the epithelium translate to changes in gut morphology are not entirely clear. For example, proliferation increases in surface fish during starvation, yet gut length decreases. The balance between cell renewal and death is likely different between the populations and in line with this, we observed that Tinaja have lower expression of tumor-suppressor and pro-apoptosis genes (i.e. *brca2, p21*-*activated kinase, caspase 6*). Overall, while we observe plasticity in gut morphology in both surface fish and cavefish, our data support the hypothesis that cavefish have a more dramatic response to nutrient fluctuations.

At post-larval stages, Pachón cavefish have altered gastrointestinal motility, resulting in delayed transit of ingested food (29). Here we found evidence of slower transit of digestive material in adult cavefish populations as well. Tinaja, Pachón, and Molino guts were full 24-hours after feeding, compared to surface fish guts that were mostly empty. Slower transit time may have evolved to optimize digestion of decaying material mixed with mud, a likely way food is obtained in the caves, and/or could result in increased nutrient absorption. Our results highlight that altered gut function and morphology likely accompany adaptation to the food sources in the cave.

We found evidence that the genetic architecture controlling gut morphology diverged between surface fish and Tinaja cavefish. We identified a QTL associated with hindgut length and found that F1 and heterozygous F2 hybrids have a longer hindgut compared to surface fish and cavefish. This phenomenon (heterosis) has been observed for another *A. mexicanus* trait, thickness of the inner retina (36), and is often seen in plant breeding, yet the genetic and molecular basis is not easy to define (37). One hypothesis is that each parent has genetic changes that result in an inferior phenotype, and that in the hybrid these changes are complemented, resulting in a superior phenotype (Dominance). Another possibility is that there are multiple alleles that interact in a way that produces a phenotype greater than what could be achieved in a homozygous condition (Over-dominance). It is possible that there are multiple alleles that have antagonistic effects on hindgut length and when they are combined the result is a longer gut.

To investigate which genes in the QTL may be important for controlling gut length we used a combination of population genetics and RNA sequencing. Our results revealed two genes that are known to regulate intestinal homeostasis in other species, *klf5* (krüppel-like factor 5) and *Notch1A*. KLF5 is a transcription factor expressed in intestinal crypts that can promote or inhibit proliferation and is commonly dysregulated in colon cancer (38–40). Notch signaling balances stem cell renewal and differentiation; activation of Notch signaling promotes stem cell proliferation and suppresses secretory cell formation (31). We found evidence that *klf5l* may be under selection in the Tinaja cave as we observed significant HapFLK p-values comparing Tinaja to Rio Choy surface fish. While we did not find the same evidence for the *notch1a* coding region, we found that expression of *notch1a* is significantly different between surface fish, Tinaja cavefish, and F1 hybrids, suggesting there may be regulatory mutations within the QTL that influence *notch1a* expression. Tinaja cavefish tend to have higher n*otch1a* expression, in line with greater proliferation in the epithelium. However, we found that notch-regulated ankyrin repeat protein (*nrarpa*) that suppresses Notch signaling in other contexts (41, 42) is more highly expressed in both Tinaja cavefish and F1 hybrids. Since Notch signaling tends to be context dependent, comparing the components of the Notch signaling pathway with cellular resolution may give insight into how differences in proliferation and morphology arise.

Cave-adapted *A. mexicanus* not only experience a different diet in the cave, but also a different microbial landscape (43). We found that complement factor B (*cfb1*) is within the hindgut QTL, shows significant divergence in all populations, and is differentially expressed in the hindgut. Complement factor B is part of the innate immune system. It is expressed in multiple tissues, including the colonic mucosa, and functions in the pathway that recognizes and eliminates bacterial pathogens by controlling immune cell differentiation (44, 45). Mounting evidences suggests that immune cells alter self-renewal of stem cells in the epithelium through cytokine signaling (5). Interestingly, we found that cavefish and F1 hybrids have lower expression of a suppressor of cytokine signaling in the hindgut. Recent research suggests that Pachón cavefish have fewer pro-inflammatory immune cells in the visceral adipose tissue (46), but the amount and differences in the immune cells in the gut have not been explored. Our results suggest that evolution of the immune system in *A. mexicanus* likely plays a role in how the gut responds to external cues. In the lab, the external microbial landscape is the same for the populations, yet it is possible that genetic differences select for specific microbes and this in turn influences intestinal homeostasis.

Cavefish accumulate more carotenoids (retinoic acid (RA) precursors) in the visceral adipose tissue compared to surface fish and this correlates with decreased expression of genes that convert carotenoids into retinoids in the gut epithelium (47). We found that cavefish and F1 hybrids have lower expression of beta-carotene oxygenase (*bco1*) and higher expression of retinol dehydrogenase (*rhd1*) in the hindgut. The outcome may be changes in RA production and signaling. RA signaling is important for stem cell maintenance in a number of contexts (48), but how it regulates intestinal stem cell homeostasis *in vivo* is not well understood. However, it is clear that RA signaling regulates innate immune cells in the gut (49). It is possible that altered RA signaling in the cavefish gut influences the crosstalk between immune cells and stem cells resulting in differences in morphology.

We found that expression of the circadian clock genes *per1* and *timeless*, which are typically synchronized, show the opposite expression pattern in the hindguts of cavefish and F1 hybrids. Proliferation and gene expression in the intestine is under circadian control and disruption of clock genes alters renewal of the epithelium (50–52). Having evolved in complete darkness, cavefish have altered circadian rhythm as evidenced by developmentally delayed and reduced amplitude of clock gene expression, lack of circadian cycles of metabolism, and reduced sleep (26, 53–55). It is plausible that alterations to the gene regulatory network controlling circadian rhythm influenced homeostasis of the intestinal epithelium during cavefish evolution. Overall, our study reveals multiple levels at which the *A. mexicanus* gut has evolved in response to ecological differences in food availability including morphology, homeostasis, and plasticity in response to dietary fluctuations. Furthermore, we implicate genetic changes and molecular pathways that may have accompanied evolution of these traits.

## Methods

### Fish husbandry and diet

Fish husbandry was performed according to (56). For fixed diet experiments fish were housed individually in 1.5L tanks and fed three pellets (approximately 6mg) of New Life Spectrum TheraA+ small fish formula once per day for at least 4 months. Commercially-available low nutrient (Hikari Goldfish Staple), moderate nutrient (New Life Spectrum TheraA+), and high-nutrient (Ken’s Premium Growth Meal #3) diets were fed *ad libitum.*

### Phenotype quantification and data visualization

Fish were euthanized in 1400ppm Tricane and dissected 24-hours post feeding. Images were taken using a Cannon Powershot D12 digital camera. ImageJ was used to measure fish length and gut length. Data visualization was performed using R (57).

### Histology and imaging

Figure 1: The fish was euthanized in 1400 ppm tricane, an incision was made along the ventral body line, and it was placed into 10% buffered formalin and shipped to FishVet Group (Portland, ME) for further processing. The gut was dehydrated using a graded ethanol series, followed by several changes in xylene, and then embedded in paraffin. Sections were taken at 5μm thickness, stained with hematoxylin and eosin (H&E) and imaged using Aperio ScanScope CS. Figure 4,5: To quantify proliferation, fish were anesthetized using 400ppm Tricane, then 40μL of EdU (500μM in .1% DMSO) was injected into the intraperitoneal cavity using an insulin syringe. After 24 hours, fish were euthanized in 1400 ppm tricane and the gut was immediately removed, incubated overnight in 10% formalin, then washed in PBS and dehydrated to 100% ethanol through a graded ethanol series. The gut was incubated briefly in Histo-clear (National Diagnostics). The midgut and hindgut were separated and embedded in paraffin side by side in a single block in anterior to posterior orientation to obtain a transverse section of each structure. Blocks were sectioned at a thickness of 5μm using a Leica RM 2155 rotary microtome. Slides were dried overnight, deparaffinized using Histo-clear, and rehydrated to water through a graded ethanol series. The slides were placed in 10% buffered formalin for 10 minutes and washed twice in 3% Bovine Serum Albumin (BSA). EdU was detected using the Click-IT EdU Cell Proliferation Kit (ThermoFisher) according to the manufacturers protocol. Slides were mounted in ProLong Gold (ThermoFisher) and images were taken using Keyence BZ2 microscope and software. We measured tissue morphology and EdU cell number manually using ImageJ (58).

### Quantitative trait loci analysis

#### F2 hybrid population

We bred a surface fish female with a Tinaja cavefish male to produce a clutch of F1 hybrids. We generated a population of surface/Tinaja F2 hybrids by interbreeding individuals from this clutch. The F2 mapping population (n=219) consisted of three clutches produced from breeding paired F1 surface/Tinaja hybrid siblings.

#### Genotype by sequencing

We extracted DNA from caudal tail fins using DNeasy Blood and Tissue DNA extraction kit (Qiagen). DNA was shipped to Novogene (Chula Vista, CA) for quality control analysis and sequencing. Samples that contained greater than 1.5 ug DNA, minimal degradation (as determined by gel electrophoresis), and OD_260_/OD_280_ ratio of 1.8 to 2.0 were used for library construction. Each genomic DNA sample (0.3∼0.6 μg) was digested with Mse1, HaeIII, and EcoR1. Digested fragments were ligated with two barcoded adapters: a compatible sticky end with the primary digestion enzyme and the Illumina P5 or P7 universal sequence. All samples were pooled and size-selected after several rounds of PCR amplification to obtain the required fragments needed to generate DNA libraries. Concentration and insert size of each library was determined using Qubit^®^ 2.0 fluorometer and Agilent^®^ 2100 bioanalyzer respectively. Finally, quantitative real-time PCR (qPCR) was used to detect the effective concentration of each library. Qualified DNA libraries had an effective concentration of greater than 2 nM and were pooled by effective concentration and expected data production. Pair-end sequencing was then performed on Illumina^®^ HiSeq platform, with the read length of 144 bp at each end. Raw Illumina genotype-by-sequencing reads were cleaned and processed through the *process_shortreads* command in the Stacks software package (59). The cleaned reads were aligned to the *Astyanax mexicanus* reference genome (AstMex102, INSDC Assembly GCA_000372685.1, Apr 2013) using the Bowtie2 software (60). The aligned reads of 4 surface fish, 4 Tinaja cavefish and 4 F_1_ surface/Tinaja hybrids were manually stacked using the *pstacks* command. We then assigned the morphotypic origin of each allele and confirmed heterozygosity in the F_1_ samples using the *cstacks* command. Finally, we used this catalog to determine the genotypes at each locus in the F2 samples with the *sstacks* and *genotypes* command. This genotype database was formatted for use in R/qtl (61)

#### Linkage map

Using R/qtl, we selected for markers that were homozygous and had opposite genotypes in cavefish versus surface fish (based on three surface and three cave individuals). A linkage map was constructed from these loci using only the F2 population. All markers that were genotyped in less than 180 individuals were omitted, as well as all individuals that had poor marker genotyping (<1500 markers). Markers that did not conform to the expected allele segregation ratio of 1:2:1 were also omitted (p < 1e^-10^). These methods produced a map with 1839 markers, 219 individuals, and 29 linkage groups. Unlinked markers and small linkage groups (<10 markers) were omitted until our map consisted of the optimal 25 linkage groups (*Astyanax mexicanus* has 2n = 50 chromosomes (62)). Each linkage group had 20-135 markers and markers were reordered by default within the linkage groups, producing a total map length of > 5000 cM. Large gaps in the map were eliminated by manually switching marker order to the best possible order within each linkage group (error.prob = 0.005). The final linkage map consisted of 1800 markers, 219 individuals and 25 linkage groups, spanning 2871 cM. Maximum spacing was 32.1 cM and average spacing was 1.6 cM.

#### QTL scan using R/qtl

Genome wide logarithm of the odds (LOD) scores were calculated using a single-QTL normal model and Haley-Knott regression. We assessed statistical significance of LOD scores by calculating the 95^th^ percentile of genome-wide maximum penalized LOD score using 1000 random permutations (scanone). We estimated the confidence intervals for identified QTL as the 1.5-LOD support interval expanded to the nearest genotyped marker (lodint).

### Population genomic metrics and analysis

We performed the following measures with GATK-processed data, including a core set of samples analyzed in previous studies (18, 28) which contained: Pachón, N = 10 (9 newly re-sequenced plus the reference reads mapped back to the reference genome); Tinaja N =10; Molino N = 9; Rascón N = 8; and Río Choy N = 9 and required six or more individuals have data for a particular site. Details of sequencing and sample processing are in (18, 28). We used VCFtools v0.1.13 (63) to calculate π, F_ST_ and *d*_XY_ and custom python scripts to calculate these metrics on a per gene basis. We identified the allele counts per population with VCFtools and used these for subsequent *d*_XY_ and fixed differences (DF) calculations. We used hapFLK v1.3 https://forge-dga.jouy.inra.fr/projects/hapflk (64) for genome-wide estimation of the hapFLK statistic of across all 44 *Astyanax mexicanus* samples and two *Astyanax aeneus* samples. For all of these metrics, we only used sites that contained six or more individuals per population and calculated the metric or included the p-values (in the case of hapFLK) for only the coding region of the gene. Candidate genes were included if (a) the comparison between Río Choy and Tinaja resulted in a maximum *d*_XY_ = 1, suggesting that there was at least one fixed difference between Río Choy and Tinaja and (b) HapFLK resulted in at least one p value less than 0.05, suggesting that the haplotype surrounding the gene exhibited evidence for non-neutral evolution.

### RNA sequencing

#### RNA extraction and cDNA synthesis

Adult *A. mexicanus* were euthanized in 1400ppm Tricane and the hindgut was immediately removed and homogenized in 0.3mL Trizol using a motorized pellet pestle, and then stored at -80°C. Total RNA was extracted using Zymo Research Direct-zol RNA MicroPrep with DNAse treatment according to the manufacturers protocol. LunaScript RT supermix kit with 1μg of RNA was used to synthesize cDNA. All samples were processed on the same day. Diluted cDNA samples (50ng/μL) were used for sequencing.

#### RNA sequencing and differential gene expression analysis

HiSeq Illumina sequencing was performed by Novogene (Chula Vista, CA). All samples were indexed and run as pools, providing an estimated 20-30 million single-end reads per sample. Samples were processed using an RNA-seq pipeline implemented in the bcbio-nextgen project (https://bcbio-nextgen.readthedocs.org/en/latest/). Raw reads were examined for quality issues using FastQC (http://www.bioinformatics.babraham.ac.uk/projects/fastqc/) to ensure library generation and sequencing are suitable for further analysis.

Reads were aligned to Ensembl build 2 of the *Astyanax mexicanus* genome, augmented with transcript information from Ensembl release 2.0.97 using STAR (65) with soft trimming enabled to remove adapter sequences, other contaminant sequences such as polyA tails and low quality sequences. Alignments were checked for evenness of coverage, rRNA content, genomic context of alignments and complexity using a combination of FastQC, Qualimap (66), MultiQC (https://github.com/ewels/MultiQC), and custom tools. Counts of reads aligning to known genes were generated by featureCounts (67) and used for further QC with the R package bcbioRNASeq [Steinbaugh MJ, Pantano L, Kirchner RD et al. bcbioRNASeq: R package for bcbio RNA-seq analysis [version 2; peer review: 1 approved, 1 approved with reservations]. F1000Research 2018, 6:1976 (https://doi.org/10.12688/f1000research.12093.2)]. In parallel, Transcripts Per Million (TPM) measurements per isoform were generated by quasialignment using Salmon (68) for downstream differential expression analysis as quantitating at the isoform level has been shown to produce more accurate results at the gene level (69). Salmon output was imported into R using tximport (70) and differential gene expression analysis and data visualization was performed using R with the DEseq 2 package (57, 71).

## Acknowledgments

Brian Martineu and Megan Peavey for fish husbandry. This work was supported by grants from the National Institutes of Health [HD089934, DK108495].

## Tables

**Table S1.**
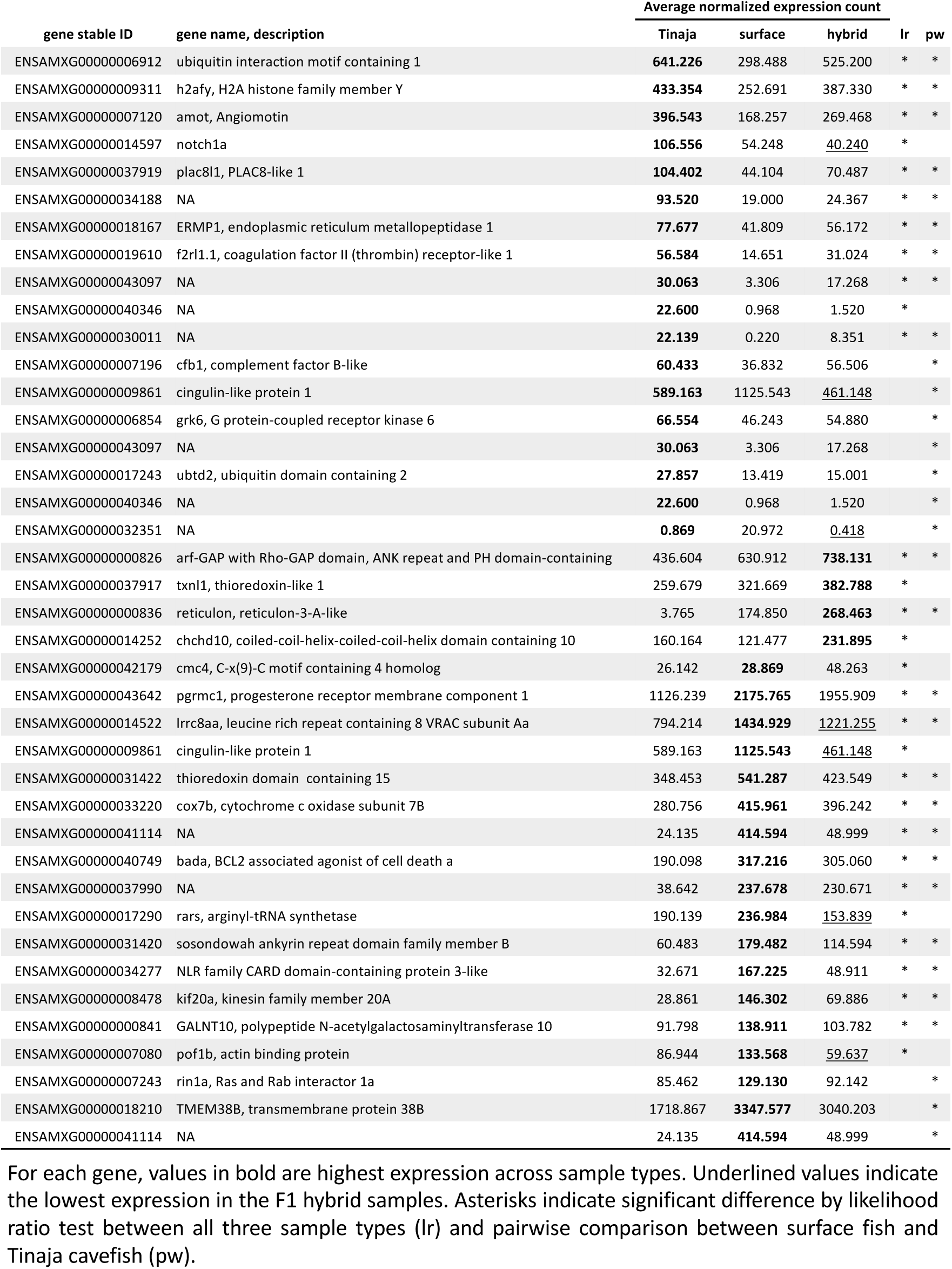
Genes within the hindgut QTL 1.5-LOD support interval that are differentially expressed in the hindgut.

**Table S2.**
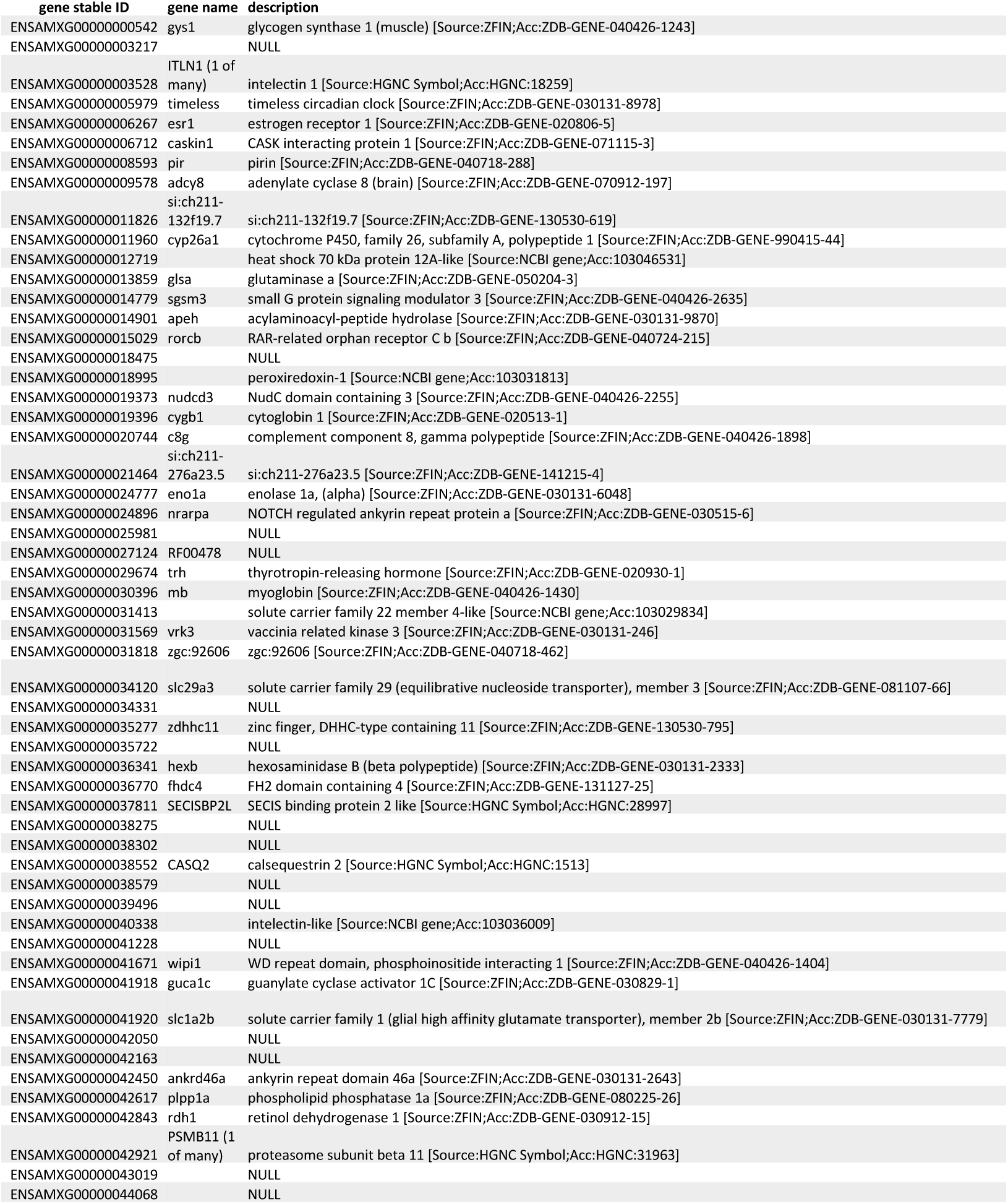
Differentially expressed hindgut genes exhibiting highest expression in the F1 hybrid (Group 3, Figure S1)

**Table S2.**
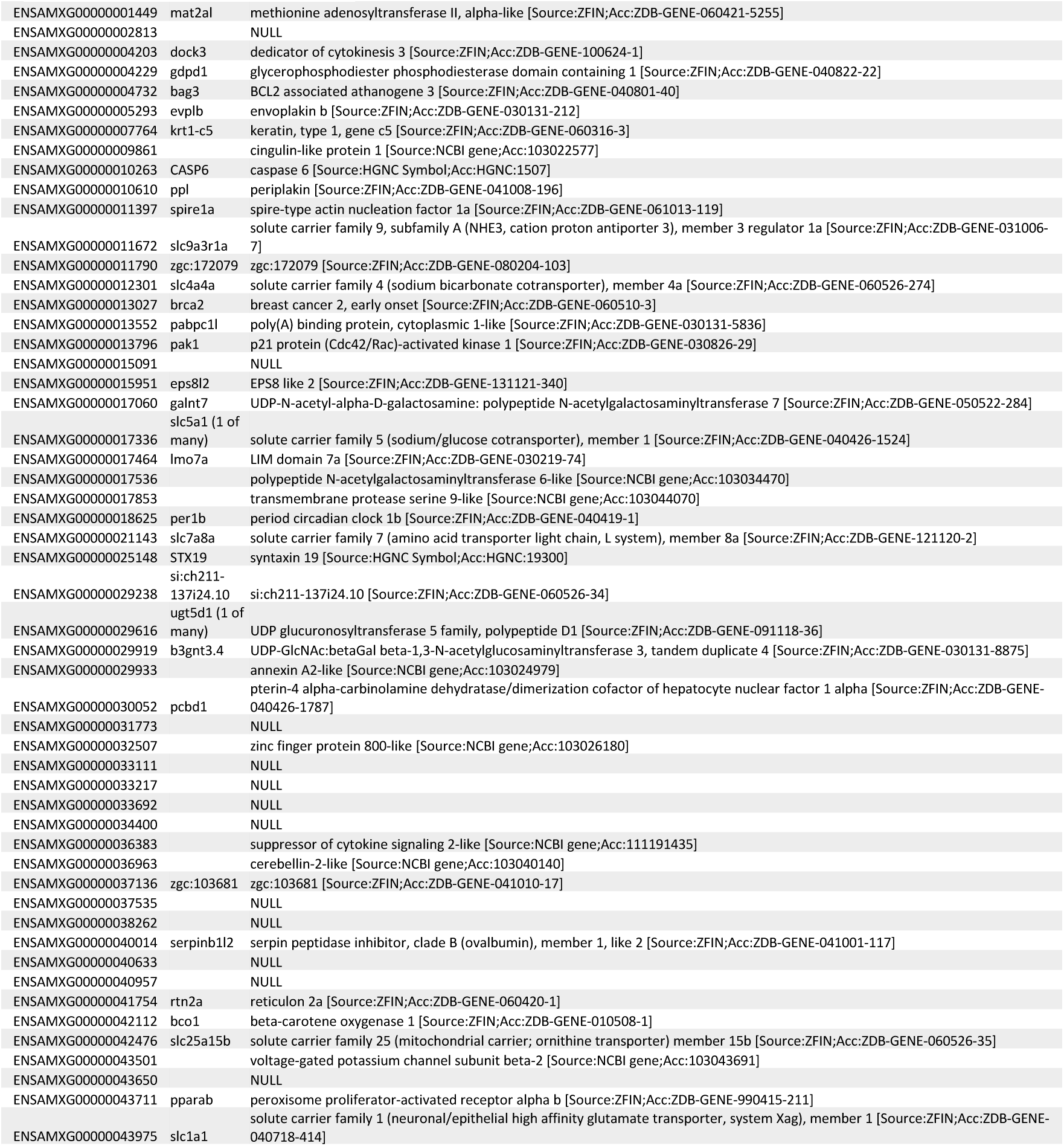
Differentially expressed hindgut genes exhibiting lowest expression in the F1 hybrid (Group 4, Figure S1)

## Figures and figure legends

**Supplemental Figure 1:**
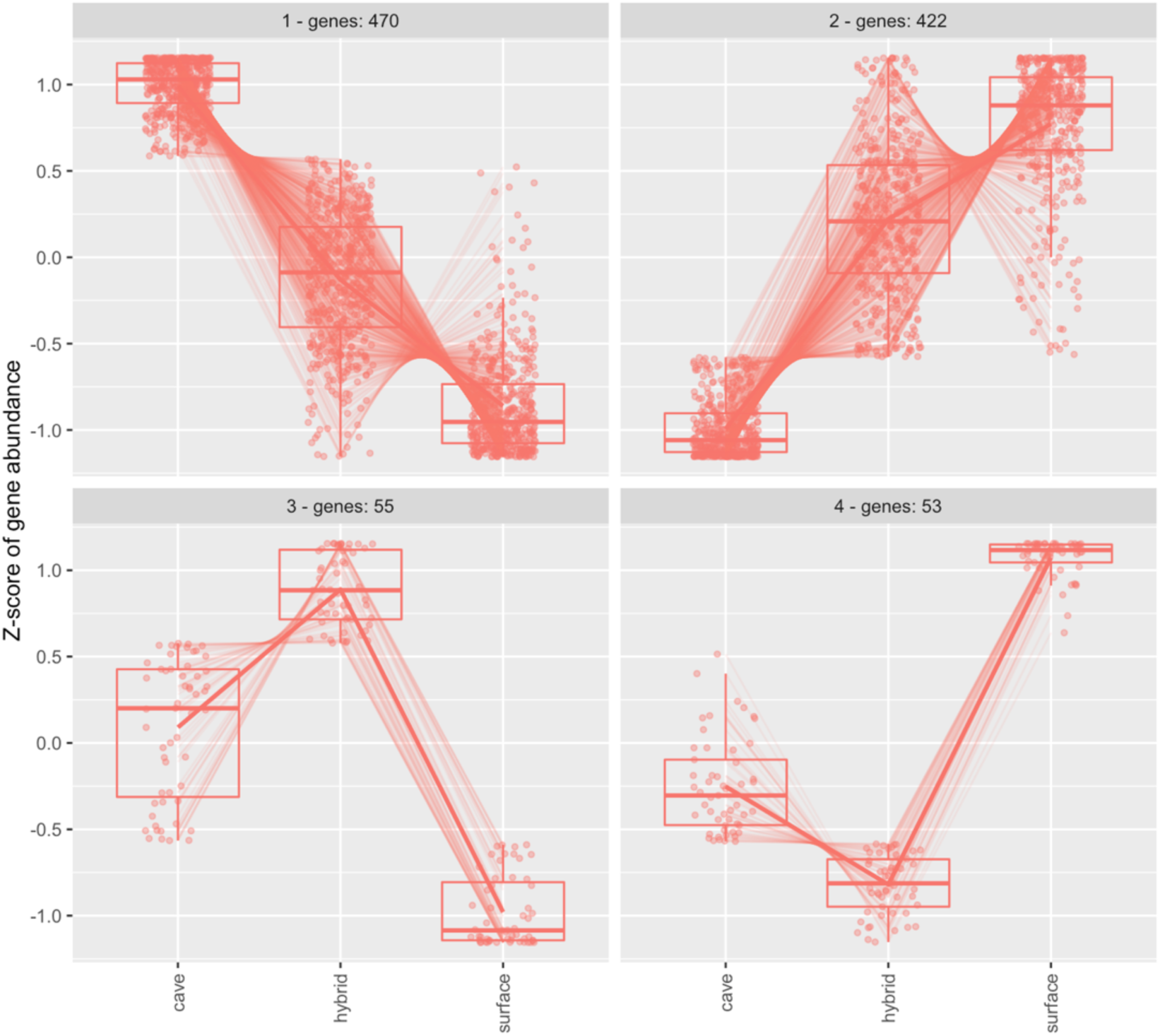
Gene expression clusters of genes that are differentially expressed between Tinaja, Surface, and F1 hybrid hindguts (n=5 hindguts per population).

## References

1. H. Gehart, H. Clevers, Tales from the crypt: new insights into intestinal stem cells. Nat. Rev. Gastroenterol. Hepatol. 16, 19–34 (2019).

2. S. Beyaz, et al., High-fat diet enhances stemness and tumorigenicity of intestinal progenitors. Nature 531, 53–58 (2016).

3. C.-W. Cheng, et al., Ketone Body Signaling Mediates Intestinal Stem Cell Homeostasis and Adaptation to Diet. Cell 178, 1115-1131.e15 (2019).

4. C. A. Lindemans, et al., Interleukin-22 promotes intestinal-stem-cell-mediated epithelial regeneration. Nature 528, 560–564 (2015).

5. M. Biton, et al., T Helper Cell Cytokines Modulate Intestinal Stem Cell Renewal and Differentiation. Cell 175, 1307-1320.e22 (2018).

6. D. M. Smith, R. C. Grasty, N. A. Theodosiou, C. J. Tabin, N. M. Nascone-Yoder, Evolutionary relationships between the amphibian, avian, and mammalian stomachs. Evol. Dev. 2, 348–359 (2000).

7. K. D. Walton, D. Mishkind, M. R. Riddle, C. J. Tabin, D. L. Gumucio, Blueprint for an intestinal villus: Species-specific assembly required. Wiley Interdiscip. Rev. Dev. Biol. 7, e317 (2018).

8. N. A. Theodosiou, E. Oppong, 3D morphological analysis of spiral intestine morphogenesis in the little skate, <scp> *Leucoraja erinacea* </scp>. Dev. Dyn. 248, 688–701 (2019).

9. S. Takashima, D. Gold, V. Hartenstein, Stem cells and lineages of the intestine: a developmental and evolutionary perspective. Dev. Genes Evol. 223, 85–102 (2013).

10. I. Miguel-Aliaga, H. Jasper, B. Lemaitre, Anatomy and Physiology of the Digestive Tract of Drosophila melanogaster. Genetics 210, 357–396 (2018).

11. N. Aghaallaei, et al., Identification, visualization and clonal analysis of intestinal stem cells in fish. Development 143, 3470–3480 (2016).

12. S. C. Leigh, B.-Q. Nguyen-Phuc, D. P. German, The effects of protein and fiber content on gut structure and function in zebrafish (Danio rerio). J. Comp. Physiol. B 188, 237– 253 (2018).

13. S. M. Secor, Digestive physiology of the Burmese python: broad regulation of integrated performance. J. Exp. Biol. 211, 3767–3774 (2008).

14. S. R. McWilliams, W. H. Karasov, Phenotypic flexibility in digestive system structure and function in migratory birds and its ecological significance. Comp. Biochem. Physiol. Part A Mol. Integr. Physiol. 128, 577–591 (2001).

15. A. L. Andrew, et al., Growth and stress response mechanisms underlying post-feeding regenerative organ growth in the Burmese python. BMC Genomics 18, 338 (2017).

16. L. Espinasa, et al., Contrasting feeding habits of post-larval and adult Astyanax cavefish. Subterr. Biol. 21, 1–17 (2017).

17. G. Spicher, [Cytochrome absorption spectra of bacteria as aid for solving taxonomic problems. 1. Elaboration and description of a method for measuring the redox and carbon monoxide difference spectra of suspensions of intact live bacterial cells (author’s transl)]. Zentralbl. Bakteriol. Orig. A. 226, 524–40 (1974).

18. A. Herman, et al., The role of gene flow in rapid and repeated evolution of cave-related traits in Mexican tetra, Astyanax mexicanus. Mol. Ecol. 27, 4397–4416 (2018).

19. M. E. Protas, et al., Genetic analysis of cavefish reveals molecular convergence in the evolution of albinism. Nat. Genet. 38, 107–111 (2006).

20. S. E. McGaugh, et al., The cavefish genome reveals candidate genes for eye loss. Nat. Commun. 5 (2014).

21. A. K. Powers, E. M. Davis, S. A. Kaplan, J. B. Gross, Cranial asymmetry arises later in the life history of the blind Mexican cavefish, Astyanax mexicanus. PLoS One 12 (2017).

22. H. Hinaux, et al., Sensory evolution in blind cavefish is driven by early embryonic events during gastrulation and neurulation. Development 143, 4521–4532 (2016).

23. J. E. Kowalko, et al., Loss of schooling behavior in cavefish through sight-dependent and sight-independent mechanisms. Curr. Biol. 23, 1874–1883 (2013).

24. M. Yoshizawa, et al., Distinct genetic architecture underlies the emergence of sleep loss and prey-seeking behavior in the Mexican cavefish. BMC Biol. 13, 15 (2015).

25. J. Jaggard, et al., The lateral line confers evolutionarily derived sleep loss in the Mexican cavefish. J. Exp. Biol. 220, 284–293 (2017).

26. D. Moran, R. Softley, E. J. Warrant, Eyeless Mexican cavefish save energy by eliminating the circadian rhythm in metabolism. PLoS One 9, e107877 (2014).

27. A. C. Aspiras, N. Rohner, B. Martineau, R. L. Borowsky, C. J. Tabin, Melanocortin 4 receptor mutations contribute to the adaptation of cavefish to nutrient-poor conditions. Proc. Natl. Acad. Sci. 112, 9668–9673 (2015).

28. M. R. Riddle, et al., Insulin resistance in cavefish as an adaptation to a nutrient-limited environment. Nature 555, 647–651 (2018).

29. M. R. Riddle, W. Boesmans, O. Caballero, Y. Kazwiny, C. J. Tabin, Morphogenesis and motility of the Astyanax mexicanus gastrointestinal tract. Dev. Biol. 441, 285–296 (2018).

30. M. R. Riddle, et al., Genetic architecture underlying changes in carotenoid accumulation during the evolution of the Blind Mexican cavefish, &lt;em&gt;Astyanax mexicanus&lt;/em&gt; bioRxiv, 788844 (2019).

31. E. S. Demitrack, L. C. Samuelson, Notch regulation of gastrointestinal stem cells. J. Physiol. 594, 4791–4803 (2016).

32. S. Xiong, J. Krishnan, R. Peuß, N. Rohner, Early adipogenesis contributes to excess fat accumulation in cave populations of Astyanax mexicanus. Dev. Biol. 441, 297–304 (2018).

33. R. K. Buddington, J. M. Diamond, Aristotle revisited: the function of pyloric caeca in fish. Proc. Natl. Acad. Sci. 83, 8012–8014 (1986).

34. N. A. Ballesteros, et al., The Pyloric Caeca Area Is a Major Site for IgM+ and IgT+ B Cell Recruitment in Response to Oral Vaccination in Rainbow Trout. PLoS One 8, e66118 (2013).

35. L. D. Townsend, Variation in the Number of Pyloric Caeca and Other Numerical Characters in Chinook Salmon and in Trout. Copeia 1944, 52 (1944).

36. K. E. O’Quin, M. Yoshizawa, P. Doshi, W. R. Jeffery, Quantitative Genetic Analysis of Retinal Degeneration in the Blind Cavefish Astyanax mexicanus. PLoS One 8, e57281 (2013).

37. J. A. Birchler, H. Yao, S. Chudalayandi, Unraveling the genetic basis of hybrid vigor. Proc. Natl. Acad. Sci. 103, 12957–12958 (2006).

38. B. B. McConnell, et al., Krüppel-Like Factor 5 Is Important for Maintenance of Crypt Architecture and Barrier Function in Mouse Intestine. Gastroenterology 141, 1302-1313.e6 (2011).

39. T. Nakaya, et al., KLF5 Regulates the Integrity and Oncogenicity of Intestinal Stem Cells. Cancer Res. 74, 2882–2891 (2014).

40. X. Zhang, et al., Somatic Superenhancer Duplications and Hotspot Mutations Lead to Oncogenic Activation of the KLF5 Transcription Factor. Cancer Discov. 8, 108–125 (2018).

41. T. Ishitani, K. Matsumoto, A. B. Chitnis, M. Itoh, Nrarp functions to modulate neural-crest-cell differentiation by regulating LEF1 protein stability. Nat. Cell Biol. 7, 1106– 1112 (2005).

42. T. J. Yun, M. J. Bevan, Notch-Regulated Ankyrin-Repeat Protein Inhibits Notch1 Signaling: Multiple Notch1 Signaling Pathways Involved In T Cell Development. J. Immunol. 170, 5834–5841 (2003).

43. P. Ornelas-García, S. Pajares, V. M. Sosa-Jiménez, S. Rétaux, R. A. Miranda-Gamboa, Microbiome differences between river-dwelling and cave-adapted populations of the fish Astyanax mexicanus (De Filippi, 1853). PeerJ 6, e5906 (2018).

44. J. Laskowski, J. M. Thurman, “Chapter 14 - Factor B” in The Complement FactsBook (Second Edition), Factsbook., Second Edi, S. Barnum, T. Schein, Eds. (Academic Press, 2018), pp. 135–146.

45. Andoh, et al., Detection of complement C3 and factor B gene expression in normal colorectal mucosa, adenomas and carcinomas. Clin. Exp. Immunol. 111, 477–483 (1998).

46. R. Peuß, et al., Single cell analysis reveals modified hematopoietic cell composition affecting inflammatory and immunopathological responses in &lt;em&gt;Astyanax mexicanus&lt;/em&gt; bioRxiv, 647255 (2019).

47. M. R. Riddle, et al., Genetic architecture underlying changes in carotenoid accumulation during the evolution of the Blind Mexican cavefish, &lt;em&gt;Astyanax mexicanus&lt;/em&gt; bioRxiv, 788844 (2019).

48. Mezquita, Mezquita, Two Opposing Faces of Retinoic Acid: Induction of Stemness or Induction of Differentiation Depending on Cell-Type. Biomolecules 9, 567 (2019).

49. P. Czarnewski, S. Das, S. Parigi, E. Villablanca, Retinoic Acid and Its Role in Modulating Intestinal Innate Immunity. Nutrients 9, 68 (2017).

50. P. Karpowicz, Y. Zhang, J. B. Hogenesch, P. Emery, N. Perrimon, The Circadian Clock Gates the Intestinal Stem Cell Regenerative State. Cell Rep. 3, 996–1004 (2013).

51. K. Parasram, P. Karpowicz, Time after time: circadian clock regulation of intestinal stem cells. Cell. Mol. Life Sci. (2019) https:/doi.org/10.1007/s00018-019-03323-x.

52. E. Peyric, H. A. Moore, D. Whitmore, Circadian Clock Regulation of the Cell Cycle in the Zebrafish Intestine. PLoS One 8, e73209 (2013).

53. A. Beale, et al., Circadian rhythms in Mexican blind cavefish Astyanax mexicanus in the lab and in the field. Nat. Commun. 4, 2769 (2013).

54. I. A. Frøland Steindal, A. D. Beale, Y. Yamamoto, D. Whitmore, Development of the Astyanax mexicanus circadian clock and non-visual light responses. Dev. Biol. 441, 345–354 (2018).

55. E. R. Duboué, A. C. Keene, R. L. Borowsky, Evolutionary convergence on sleep loss in cavefish populations. Curr. Biol. 21, 671–676 (2011).

56. Y. Elipot, L. Legendre, S. Père, F. Sohm, S. Rétaux, *Astyanax* Transgenesis and Husbandry: How Cavefish Enters the Laboratory. Zebrafish 11, 291–299 (2014).

57. R-core-team, R: A Language and Environment for Statistical Computing (2019).

58. C. T. Rueden, et al., ImageJ2: ImageJ for the next generation of scientific image data. BMC Bioinformatics 18 (2017).

59. J. M. Catchen, A. Amores, P. Hohenlohe, W. Cresko, J. H. Postlethwait, Stacks: building and genotyping Loci de novo from short-read sequences. G3 (Bethesda). 1, 171–82 (2011).

60. B. Langmead, S. L. Salzberg, Fast gapped-read alignment with Bowtie 2. Nat. Methods 9, 357–9 (2012).

61. D. Arends, P. Prins, R. C. Jansen, K. W. Broman, R/qtl: high-throughput multiple QTL mapping: Fig. 1. Bioinformatics 26, 2990–2992 (2010).

62. K. F. Kavalco, L. F. De Almeida-Toledo, Molecular Cytogenetics of Blind Mexican Tetra and Comments on the Karyotypic Characteristics of Genus Astyanax (Teleostei, Characidae). Zebrafish 4, 103–111 (2007).

63. P. Danecek, et al., The variant call format and VCFtools. Bioinformatics 27, 2156– 2158 (2011).

64. M. I. Fariello, S. Boitard, H. Naya, M. SanCristobal, B. Servin, Detecting Signatures of Selection Through Haplotype Differentiation Among Hierarchically Structured Populations. Genetics 193, 929–941 (2013).

65. A. Dobin, et al., STAR: ultrafast universal RNA-seq aligner. Bioinformatics 29, 15–21 (2013).

66. F. García-Alcalde, et al., Qualimap: evaluating next-generation sequencing alignment data. Bioinformatics 28, 2678–2679 (2012).

67. Y. Liao, G. K. Smyth, W. Shi, featureCounts: an efficient general purpose program for assigning sequence reads to genomic features. Bioinformatics 30, 923–930 (2014).

68. N. Bray, H. Pimentel, P. Melsted, L. Pachter, Near-optimal RNA-Seq quantification. bioRxiv (2015) https:/doi.org/10.1101/021592.

69. C. Soneson, M. I. Love, M. D. Robinson, Differential analyses for RNA-seq: transcript-level estimates improve gene-level inferences. F1000Research 4, 1521 (2016).

70. C. Soneson, M. I. Love, M. D. Robinson, Differential analyses for RNA-seq: transcript-level estimates improve gene-level inferences. F1000Research 4, 1521 (2015).

71. M. I. Love, W. Huber, S. Anders, Moderated estimation of fold change and dispersion for RNA-seq data with DESeq2. Genome Biol. 15, 550 (2014).

